# Microglia Regulate Sleep via Calcium-Dependent Modulation of Norepinephrine Transmission

**DOI:** 10.1101/2023.07.24.550176

**Authors:** Chenyan Ma, Bing Li, Daniel Silverman, Xinlu Ding, Anan Li, Chi Xiao, Ganghua Huang, Kurtresha Worden, Sandra Muroy, Wei Chen, Zhengchao Xu, Chak Foon Tso, Yixuan Huang, Yufan Zhang, Qingming Luo, Kaoru Saijo, Yang Dan

**Author notes:** Dascena, 12333 Sowden Rd Ste B, Houston, TX 77080, USA. These authors contributed equally to this work (Chenyan Ma, Bing Li).

## Abstract

Sleep interacts reciprocally with immune system activity, but its specific relationship with microglia – the resident immune cells in the brain – remains poorly understood. Here we show that microglia can regulate sleep through a mechanism involving Gi-coupled GPCRs, intracellular Ca^2+^ signaling, and suppression of norepinephrine transmission. Chemogenetic activation of microglia Gi signaling strongly promoted sleep, whereas pharmacological blockade of Gi-coupled P2Y12 receptors decreased sleep. Two-photon imaging showed that P2Y12/Gi activation elevated microglia intracellular Ca^2+^, and blockade of this Ca^2+^ elevation largely abolished the Gi-induced sleep increase. Microglia Ca^2+^ level also increased at natural wake-to-sleep transitions, caused partly by reduced norepinephrine. Furthermore, imaging of norepinephrine activity with its biosensor showed that microglia P2Y12/Gi activation significantly reduced norepinephrine, partly by increasing the adenosine concentration. Thus, microglia can regulate sleep through reciprocal interactions with norepinephrine transmission.

## Main

Sleep plays a vital role in brain health and function by facilitating multiple physiological processes, including homeostatic regulation of neuronal activity, synaptic strengths, and clearance of metabolic waste products^1-4^. Microglia, the primary immune cells in the brain, play a key part in brain homeostasis by modulating neuronal activity, pruning synapses, and clearing cellular debris and harmful aggregates^5-11^. Both sleep disturbances and microglia dysfunction have been implicated in multiple neurodegenerative diseases^5, 7, 12–14^. However, the role of microglia in sleep regulation is only beginning to be investigated^15–17^. In addition to specific neuronal circuits controlling sleep^18^, several metabolic substances (e.g., ATP and adenosine) and immune-modulating cytokines (e.g., IL-1β and TNFα) have been shown to promote sleep^19^. Microglia are well-suited for mediating such sleep-regulating effects, as they constantly survey the brain parenchyma with their motile processes, both sensing and responding to purinergic molecules and cytokines^5, 8, 9^.

In a healthy brain, microglia exist in a homeostatic state, characterized by ramified morphology and expression of specific genes supporting homeostatic functions^6, 9^. One of these homeostatic genes is P2Y12, a Gi-protein coupled ATP/ADP receptor that is expressed specifically in microglia within the central nervous system. P2Y12 is crucial for microglia function, particularly in their sensing and modulation of neuronal activity^20–23^, facilitation of experience-dependent plasticity^24^, and protection against epilepsy^20^ and ischemia-induced brain injury^22^. The ligands for P2Y12 – ATP and ADP – as well as their metabolite adenosine are known to play important roles in homeostatic sleep regulation^25^.

We thus set out to study the role of microglia P2Y12/Gi signaling in regulating sleep. Using *Tmem119-CreERT2* mice for microglia-specific imaging and manipulation, we found that microglia P2Y12/Gi activation promoted sleep through a mechanism that depended on their intracellular Ca^2+^ signaling. Microglia Ca^2+^ activity was naturally higher during sleep than wakefulness, caused at least in part by a lower level of norepinephrine (NE). Conversely, microglia P2Y12/Gi activation reduced NE transmission, partly by increasing the level of extracellular adenosine.

## Results

### Activation of microglia Gi signaling promotes sleep

To test whether activation of Gi signaling in microglia affects sleep, we crossed *Tmem119-CreERT2* driver mice^26^ with reporter mice carrying Cre-inducible hM4Di Gi-DREADD (designer receptor exclusively activated by designer drugs), which allowed specific expression of hM4Di in microglia (Fig. 1a-c; Extended Data Fig. 1). Sleep-wake states were measured in freely moving mice in their home cage, and wake and sleep states were classified based on electroencephalogram (EEG) and electromyogram (EMG) recordings. Compared to the control experiment with saline injection, chemogenetic activation of microglia Gi signaling induced by intraperitoneal (i.p.) injection of clozapine-N-oxide (CNO, 1 mg/kg) caused a significant increase in non-rapid eye movement (NREM) sleep and decrease in wakefulness during both light and dark phases (Fig. 1d, e; Extended Data Fig. 2a, b), primarily due to an increase in the mean duration of NREM sleep episodes (Fig. 1f, g; Extended Data Fig. 2c-f). In control mice expressing CreERT2 but not Gi-DREADD, CNO had no significant effect (Extended Data Fig. 2g, h).

**Fig. 1.**
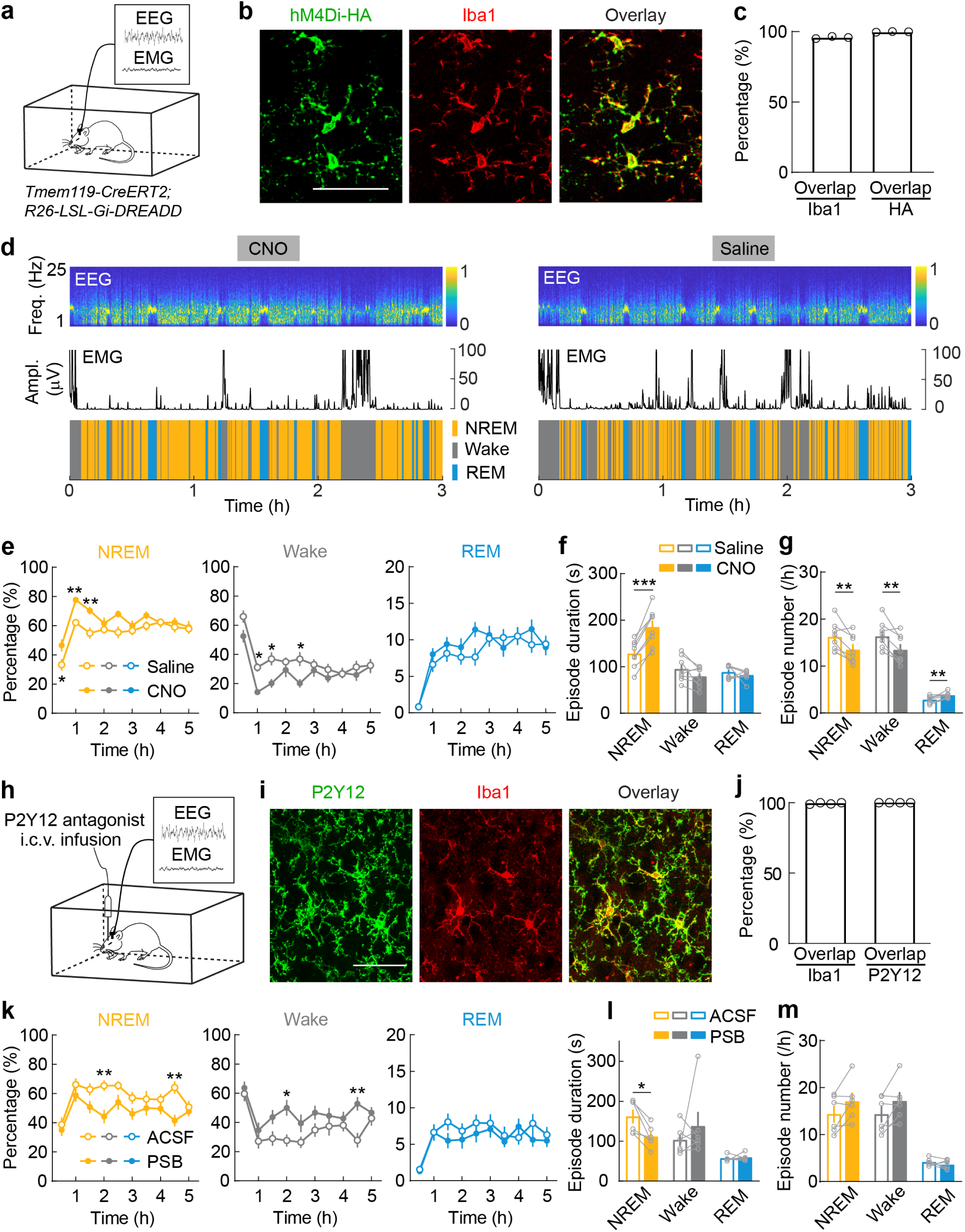
Microglia regulate sleep through P2Y12/Gi signaling. **a**, Schematic for chemogenetic experiment. **b**, Confocal images from the prefrontal cortex showing hM4Di (Gi-DREADD) expression (detected by an HA-tag antibody) in Iba1+ microglia. **c**, Quantification of efficiency and specificity (n = 3 mice, including all brain regions shown in Extended Data Fig.1). Scale bar, 50 μm. **d**, Example chemogenetic experiments with CNO or saline injection. Shown are EEG spectrogram (Freq., frequency), EMG amplitude (Ampl.), and brain states (color coded). **e**, Summary of chemogenetic experiments. Shown are the percentages of time in each brain state following CNO and saline injection (mean ± s.e.m; n = 8 mice). \**P* < 0.05, \*\**P* < 0.01 (two-way ANOVA with Bonferroni correction; NREM: treatment, *P* < 0.0001; time, *P* < 0.0001; Wake: treatment, *P* < 0.0001; time, *P* < 0.0001; REM: treatment, *P* = 0.13; time, *P* < 0.0001). **f**, **g**, Mean episode duration (**f**) and episode number per hour (**g**) for each brain state within 3 h after CNO or saline injection. Each circle indicates data from one mouse (mean ± s.e.m; n = 8 mice). \*\**P* < 0.01, \*\*\**P* < 0.001 (paired two-tailed *t*-test). **h**, Schematic for P2Y12 antagonist infusion. i.c.v., intracerebroventricular. **i**, **j**, Confocal images (**i**) and quantification (**j**) of P2Y12 expression in Iba1+ microglia in the prefrontal cortex (n = 4 mice). Scale bar, 50 μm. ‘Overlap’ refers to cells expressing both Iba1 and P2Y12. **k**, Percentage of time in each brain state following PSB0739 (PSB) and ACSF infusion (mean ± s.e.m; n = 6 mice). \**P* < 0.05, \*\**P* < 0.01 (two-way ANOVA with Bonferroni correction; NREM: treatment, *P* = 0.0005; time, *P* < 0.0001; Wake: treatment, *P* = 0.0011; time, *P* < 0.0001; REM: treatment, *P* = 0.23; time, *P* < 0.0001). **l**, **m**, Mean episode duration (**l**) and episode number per hour (**m**) for each brain state within 5 h after PSB or ACSF infusion. Each circle indicates data from one mouse (mean ± s.e.m; n = 6 mice). \**P* < 0.05 (Wilcoxon signed rank test).

**Fig. 2.**
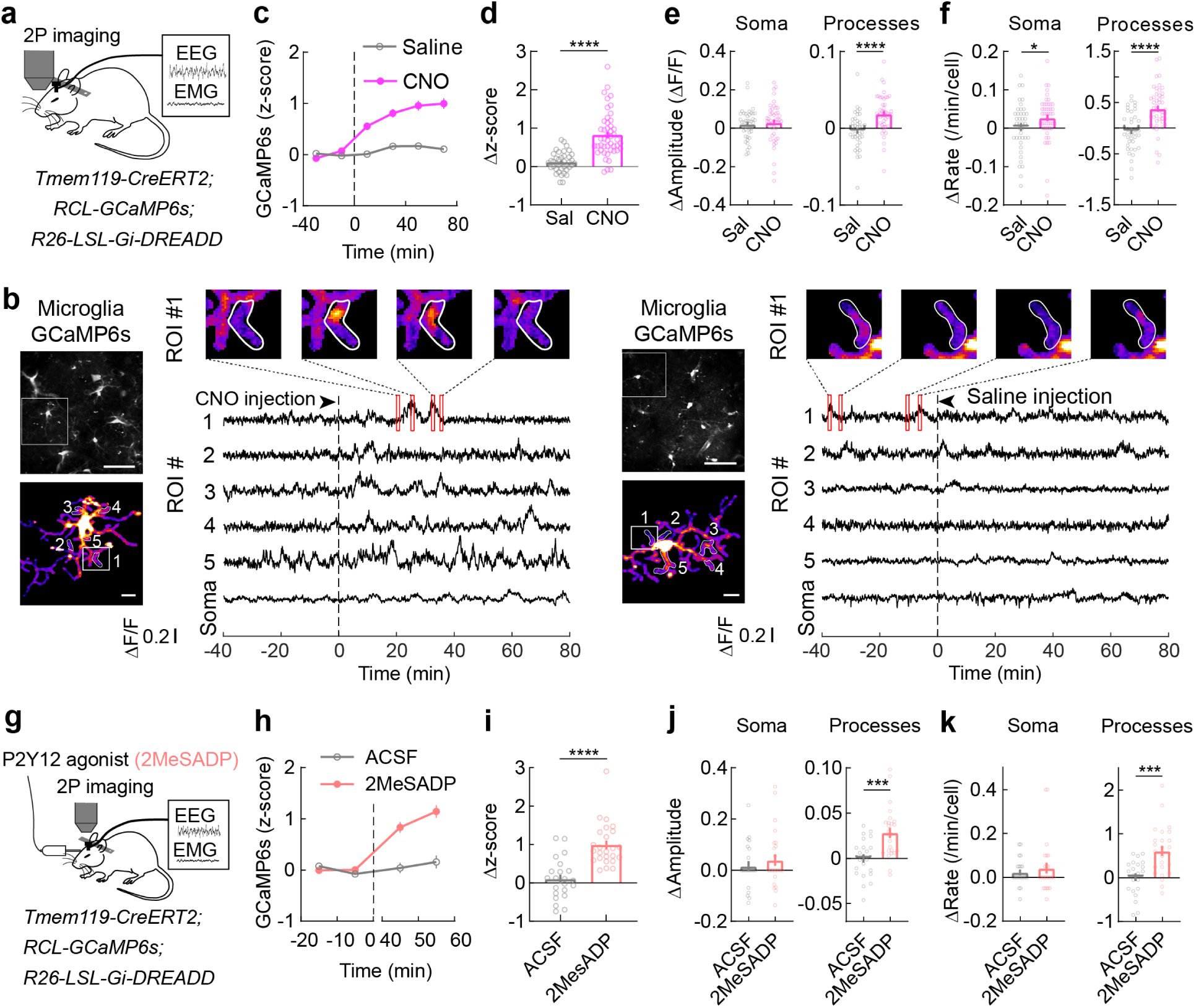
Activation of microglia Gi signaling increases intracellular Ca^2+^. **a**, Schematic for two-photon (2P) Ca^2+^ imaging in head-fixed mice. **b**, Example imaging sessions with CNO and saline injection. Top left, field of view (scale bar, 50 μm); bottom left, high magnification view of the microglia soma and processes in the white box (scale bar, 10 μm); 5 ROIs in microglia processes are outlined, whose Ca^2+^ traces are shown on the right, with snap shots of Ca^2+^ transients shown on top. Dashed line indicates time of CNO or saline injection. **c**, Z-scored Ca^2+^ activity averaged across all microglia (CNO: n = 47; saline: n = 40; from 5 mice). Dashed line, time of injection. **d-f**, Population summary of CNO-induced change in mean Ca^2+^ level (**d**), amplitude (**e**) and frequency (**f**) of Ca^2+^ events in soma and processes (difference between before and after injection). Each circle indicates data from one cell. Bars, mean ± s.e.m; \**P* < 0.05, \*\*\*\**P* < 0.0001 (Mann-Whitney *U* test). **g**, Schematic for 2P Ca^2+^ imaging with local application of P2Y12 agonist (2MeSADP). **h-k**, Similar to (**c-f**), but for local infusion of 2MeSADP or ACSF (2MeSADP: n = 25; ACSF: n = 22; from 5 mice). Dashed line, time of drug application. Each circle indicates data from one cell. Bars, mean ± s.e.m; \*\*\**P* < 0.001, \*\*\*\**P* < 0.0001 (unpaired two-tailed *t*-test or Mann-Whitney *U* test).

Next, to test whether endogenous Gi signaling mediated by P2Y12 receptors plays a role in sleep-wake regulation, we performed intracerebroventricular (i.c.v.) infusion of PSB0739 (1mM, 2 μl), a selective P2Y12 receptor antagonist^22^ (Fig. 1h-j). Compared to the control experiment with artificial cerebrospinal fluid (ACSF) infusion, PSB0739 caused a significant decrease in NREM sleep and increase in wakefulness, due mainly to a decrease in NREM episode duration (Fig. 1k-m). In contrast, activation of P2Y12 receptors with i.c.v. infusion of their agonist 2MeSADP (300 mM, 2μl) increased the episode duration of NREM sleep (Extended Data Fig. 2i-k). Thus, P2Y12 receptor signaling contributes significantly to NREM sleep, especially its maintenance.

### Gi activation increases microglia Ca^2+^ activity

In addition to a decrease in cAMP that is expected from the canonical pathway, Gi activation can also cause an increase in intracellular Ca^2+^ in microglia *in vitro*^27^. Although Ca^2+^ signaling has been shown to be involved in P2Y12-mediated chemotaxis *in vitro*^28^, its function *in vivo* is only beginning to be investigated^29, 30^.

To measure Ca^2+^ activity following Gi-DREADD activation, we performed two-photon imaging in the prefrontal cortex of mice expressing both Gi-DREADD and GCaMP6s specifically in microglia (*Tmem119-CreERT2; RCL-GcaMP6s; R26-LSL-Gi-DREADD*) (Fig. 2a, b). Under the baseline condition, we observed infrequent Ca^2+^ transients (Fig. 2b), consistent with previous studies^29, 30^. However, CNO-induced Gi activation caused a strong increase in Ca^2+^ activity, primarily in microglia processes. As shown in the population average of GCaMP6s fluorescence, CNO injection caused a progressive Ca^2+^ increase over a period of tens of minutes, whereas saline injection had no significant effect (Fig. 2c; Supplementary Video 1). The increase in Ca^2+^ induced by i.p. injection was significantly higher for CNO than saline for mice expressing Gi-DREADD in microglia (Fig. 2d, *P* < 0.0001), but not in control mice without Gi-DREADD (Extended Data Fig. 3a-c, *P* = 0.73). Note, however, that CNO-induced Gi activation also increased NREM sleep (Fig. 1a-g), which could in principle contribute to microglia Ca^2+^ increase separately from the direct effect of Gi activation. When we compared Ca^2+^ activity before and after i.p. injection within the same brain state (NREM before with NREM after, wake before with wake after), the effect of CNO was still highly significant (Extended Data Fig. 3d, e), indicating that the Ca^2+^ increase was not merely an indirect effect of brain state changes. In addition to the overall fluorescence, we also used a threshold-based method to detect discrete Ca^2+^ events and quantified their amplitudes and frequency^30^. Compared to saline control, CNO injection caused significant increases in both the amplitude and frequency of Ca^2+^ events, with stronger effects in microglia processes than the soma (Fig. 2e, f; Extended Data Fig. 3f, g). Besides chemogenetic activation of Gi signaling, local application of the P2Y12 agonist 2MeSADP (10 μM, 2 μl) in the prefrontal cortex caused a similar increase in microglia Ca^2+^ activity (Fig. 2g-k).

**Fig. 3.**
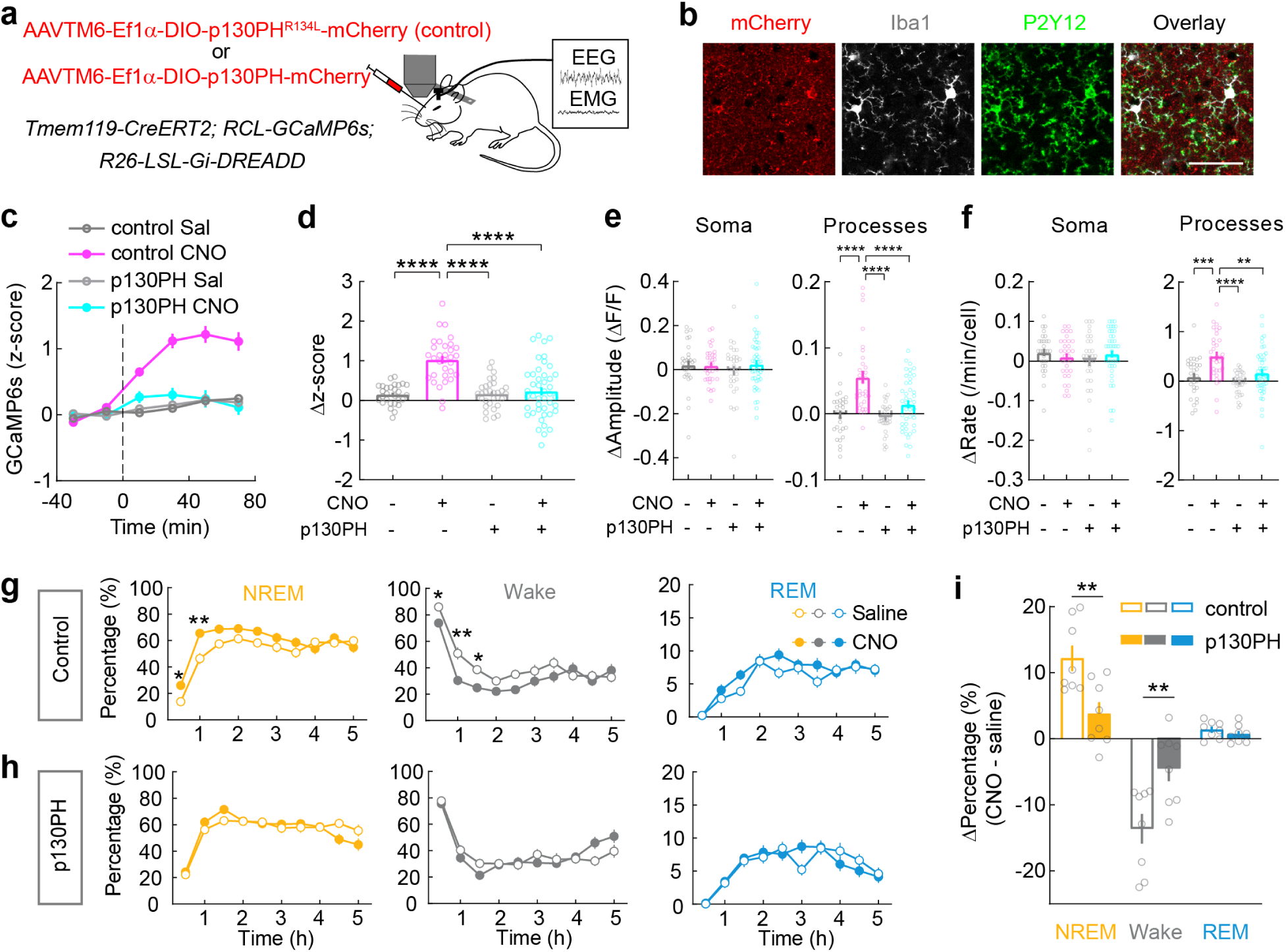
Effect of microglia Gi activation on sleep depends on increased intracellular Ca^2+^. **a**, Schematic for 2P Ca^2+^ imaging with p130PH or p130PH^R134L^ expression. **b**, Confocal images of p130PH-mCherry expression in Iba1+, P2Y12+ microglia in the prefrontal cortex. Scale bar, 50 μm. **c**, Z-scored Ca^2+^ activity averaged across all microglia imaged from mice expressing p130PH (CNO: n = 44; saline: n = 31; from 5 mice) or p130PH^R134L^ (CNO: n = 32; saline: n = 29; from 5 mice). Dashed line, time of injection. **d-f**, Population summary of CNO-induced change in mean Ca^2+^ level (**d**), amplitude (**e**) and frequency (**f**) of Ca^2+^ events in soma and processes (difference between before and after injection) in mice expressing p130PH or p130PH^R134L^. Each circle indicates data from one cell. Bars, mean ± s.e.m; \*\**P* < 0.01, \*\*\*\**P* < 0.0001 (One-way ANOVA with Holm-Šídák’s test; **d**, *P* < 0.0001; **e**, left, *P* = 0.65; right, *P* < 0.0001; **f**, left, *P* = 0.50; right, *P* < 0.0001). **g**, **h**, Effect of microglia Gi activation on sleep in mice expressing p130PH^R134L^ (**g**, n = 8 mice) or p130PH (**h**, n = 8 mice). \**P* < 0.05, \*\**P* < 0.01 (two-way ANOVA with Bonferroni correction; p130PH^R134L^, NREM: treatment, *P* < 0.0001; time, *P* < 0.0001; Wake: treatment, *P* = 0.0002; time, *P* < 0.0001; REM: treatment, *P* = 0.036; time, *P* < 0.0001; p130PH, NREM: treatment, *P* = 0.94; time, *P* < 0.0001; Wake: treatment, *P* = 0.95; time, *P* < 0.0001; REM: treatment, *P* = 0.97; time, *P* < 0.0001). **i**, Changes in each brain state induced by chemogenetic activation (difference between CNO and saline injections, averaged across 3-h after injection) in mice expressing p130PH^R134L^ or p130PH. Each circle indicates data from one mouse (mean ± s.e.m). \*\**P* < 0.01 (unpaired two tailed *t*-test).

### Role of microglia Ca^2+^ in sleep regulation

We next tested the causal role of microglia Ca^2+^ activity in regulating sleep. A previous *in vitro* study implicated the phospholipase C (PLC)-inositol trisphosphate (IP3)-Ca^2+^ cascade in P2Y12-mediated microglia chemotaxis^28^. We expressed the Pleckstrin Homology domain of PLC-like protein p130 (p130PH), which buffers cytosolic IP3 to inhibit Ca^2+^ release from the internal store^31, 32^. AAVTM6 (AAV6 with a triple mutation that transduces microglia more efficiently^33^) with Cre-dependent expression of either p130PH or p130PH^R134L^ (a mutated form of p130PH that does not bind to IP3) was injected into *Tmem119-CreERT2; RCL-GCaMP6s; R26-LSL-Gi-DREADD* mice (Fig. 3a, b; Extended Data Fig. 4). While in control mice (expressing p130PH^R134L^) CNO-induced Gi activation caused a significant Ca^2+^ elevation, no significant change was observed in mice expressing p130PH; the effect of Gi activation on Ca^2+^ activity was significantly different between p130PH and p130PH^R134L^ mice (Fig. 3c-f). Importantly, the effect of Gi activation on sleep was also largely abolished in p130PH but not p130PH^R134L^ mice (Fig. 3g-i; Extended Data Fig. 5a-d). This suggests that microglia Ca^2+^ activity is necessary for the sleep-promoting effect.

**Fig. 4.**
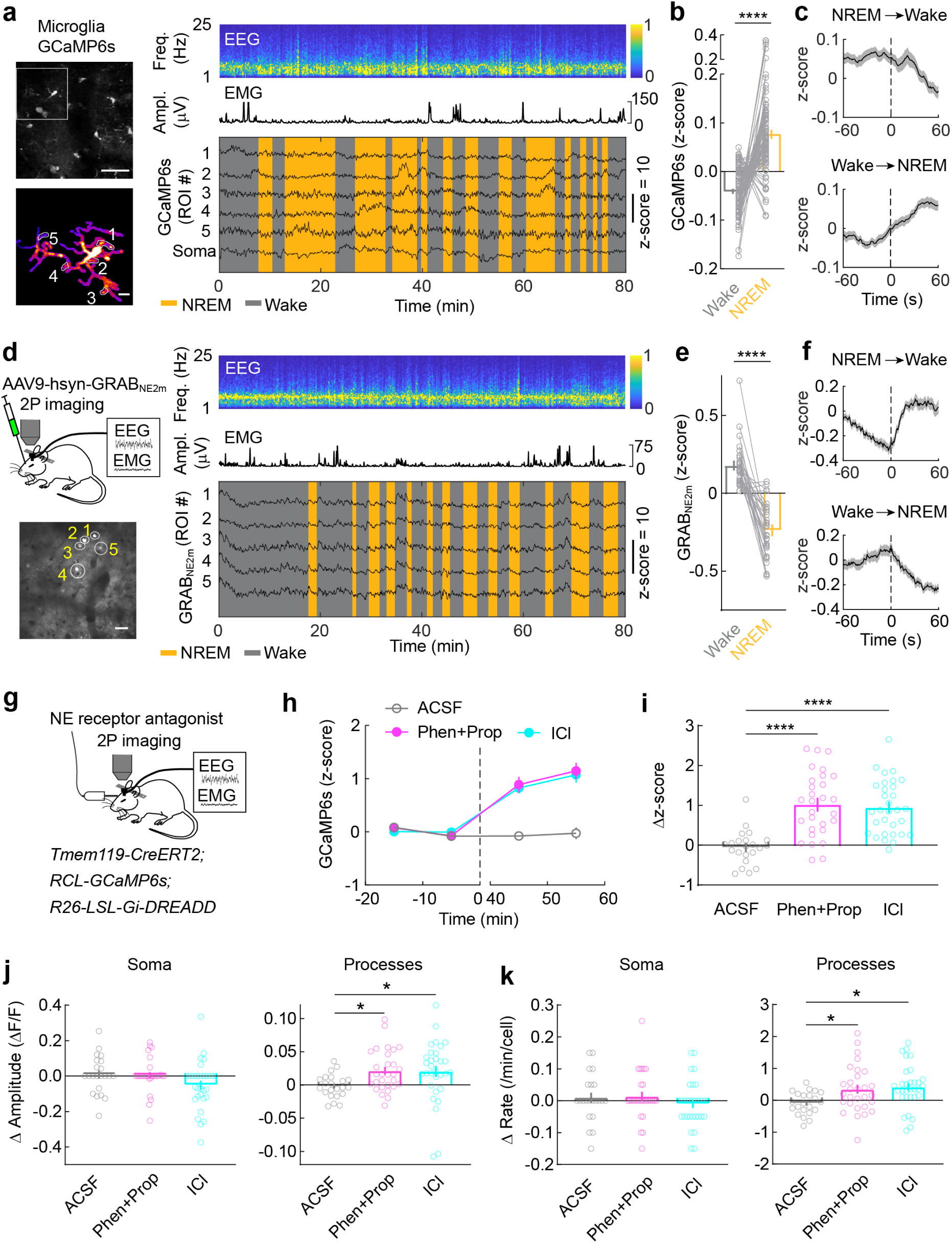
Modulation of microglia Ca^2+^ by brain state and NE. **a**, An example Ca^2+^ imaging session. Top left, field of view containing multiple microglia (scale bar, 50 μm); Bottom left, high magnification view of the microglia soma and processes in the white box (scale bar, 10 μm); 5 ROIs in processes are outlined, whose Ca^2+^ traces are shown on the right together with EEG spectrogram (Freq., frequency), EMG amplitude (Ampl.), and color-coded brain states. **b**, Summary of microglia Ca^2+^ activity during wake and NREM states. Each line presents data from one cell (n = 87 cells, from 5 mice). Bars, mean ± s.e.m.; \*\*\*\**P* < 0.0001 (Wilcoxon signed rank test). **c**, Ca^2+^ activity at brain state transitions averaged across the 87 microglia. Dashed line indicates time of transition; shading, ± s.e.m. **d**, Imaging of NE signals. Top left, schematic for 2P imaging of GRAB_NE2m_ fluorescence in the prefrontal cortex; bottom left, field of view of an example session (scale bar, 50 μm); right, NE traces of ROIs indicated in bottom left image. **e**, Average NE signals in wake and NREM states. Each line represents data from one session (n = 30 sessions, from 14 mice). Bars, mean ± s.e.m.; \*\*\*\**P* < 0.0001 (Wilcoxon signed rank test). **f**, NE signals at brain state transitions averaged across 30 sessions. Dashed line indicates time of transition; shading, ± s.e.m. **g**, Schematic of microglia Ca^2+^ imaging with local application of NE receptor antagonist. **h**, microglia Ca^2+^ before and after application of ICl (β2 receptor antagonist), Phen (α receptor antagonist) and Prop (β receptor antagonist), or ACSF, averaged across 31 cells (ICl), 29 cells (Phen+Prop), or 22 cells (ACSF) from 5 mice. Dashed line indicates time of drug application. **i-k**, Difference in mean Ca^2+^ level (**i**), amplitude (**j**) and frequency (**k**) of Ca^2+^ events before and after drug application. Each circle indicates data from one cell. Bars: mean ± s.e.m.; \**P* < 0.05, \*\*\*\**P* < 0.0001 (One-way ANOVA with Holm-Šídák’s test; **i**, *P* < 0.0001; **j**, left, *P* = 0.10; right, *P* = 0.045; **k**, left, *P* = 0.44; right, *P* = 0.014).

**Fig. 5.**
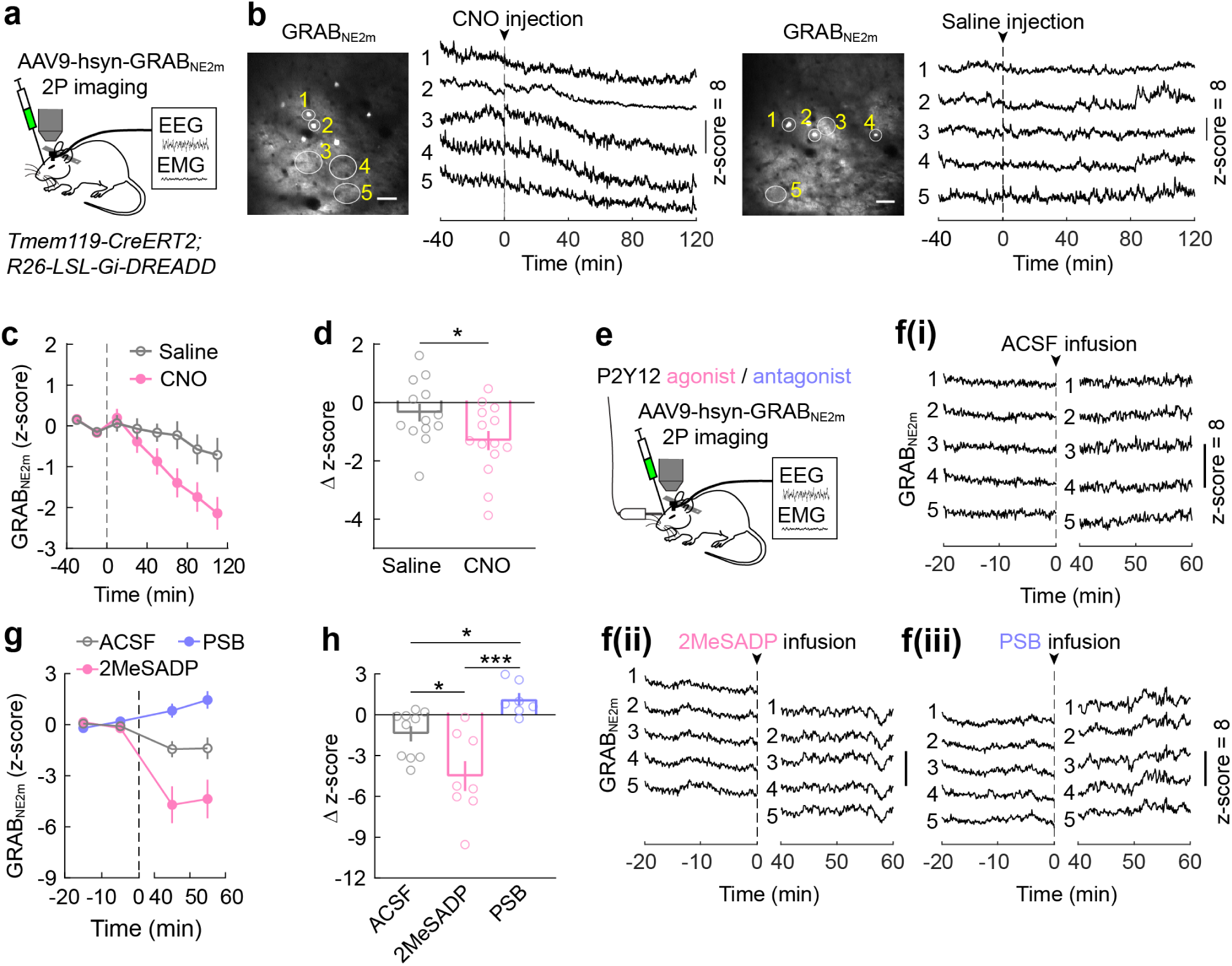
Activation of microglia Gi signaling suppresses NE transmission. **a**, Schematic of GRAB_N__E2m_ imaging in the prefrontal cortex. **b**, Example imaging session with chemogenetic Gi activation in microglia. Left, field of view (scale bar, 50 μm); 5 ROIs are outlined whose NE traces are shown on the right. Dashed line, time of CNO or saline injection. **c**, Effect of chemogenetic Gi activation in microglia on NE signals averaged across 13 (saline) or 14 (CNO) sessions from 5 mice. Dashed line indicates time of injection. **d**, Difference in NE before and after (20 – 120 min) saline or CNO injection. Each circle represents data from one session. Bars, mean ± s.e.m.; \**P* < 0.05 (unpaired two tailed *t*-test). **e**, Schematic of GRAB_NE2m_ imaging with local application of P2Y12 agonist or antagonist. **f(ⅰ)-f(ⅲ)**, NE traces from example imaging sessions with ACSF (**f(ⅰ)**), 2MeSADP (**f(ⅱ)**), or PSB (**f(ⅲ)**) application. Dashed line indicates the time of drug application. **g**, **h**, Similar to (**c**, **d**) for local drug application experiments (**e**); 2MeSADP, n = 8 sessions; PSB, n = 7; ACSF, n = 10, from 4 mice. Dashed line indicates time of drug application. \**P* < 0.05, \*\*\**P* < 0.001 (One-way ANOVA with Holm-Šídák’s test, *P* = 0.0002).

In addition to suppressing the Gi-induced increase in Ca^2+^ activity, we also elevated microglia Ca^2+^ through Gq-mediated activation of PLC-IP3 signaling. In mice expressing hM3Dq (Gq-DREADD) specifically in microglia (*Tmem119-CreERT2; R26-LSL-Gq-DREADD*), CNO-induced Gq activation significantly increased NREM sleep, with a magnitude comparable to that caused by Gi activation (Extended Data Fig. 5e-j). Together, these results indicate that microglia Ca^2+^ activity plays an important role in sleep regulation.

### Microglia Ca^2+^ activity across natural sleep-wake states

We next examined whether microglia Ca^2+^ levels naturally change between sleep and wake states. After extensive habituation, mice exhibited multiple episodes of wakefulness, NREM and REM sleep during each imaging session under the head-fixed condition^34^ (Extended Data Fig. 6a). Since during REM sleep, fluorescence imaging may be confounded by non-Ca^2+^-related changes such as in metabolic rate and blood flow^35^ (Extended Data Fig. 6b), we focused on NREM sleep and wakeful states. We observed a significant decrease in microglia Ca^2+^ at NREM→wake transitions and an increase at wake→NREM transitions, resulting in a higher level of Ca^2+^ activity during NREM sleep than wakefulness (Fig. 4a-c). Together with the observation that Gi-induced Ca^2+^ increase promoted NREM sleep, this suggests that endogenous microglia Ca^2+^ activity both regulates and is regulated by sleep-wake states.

**Fig. 6.**
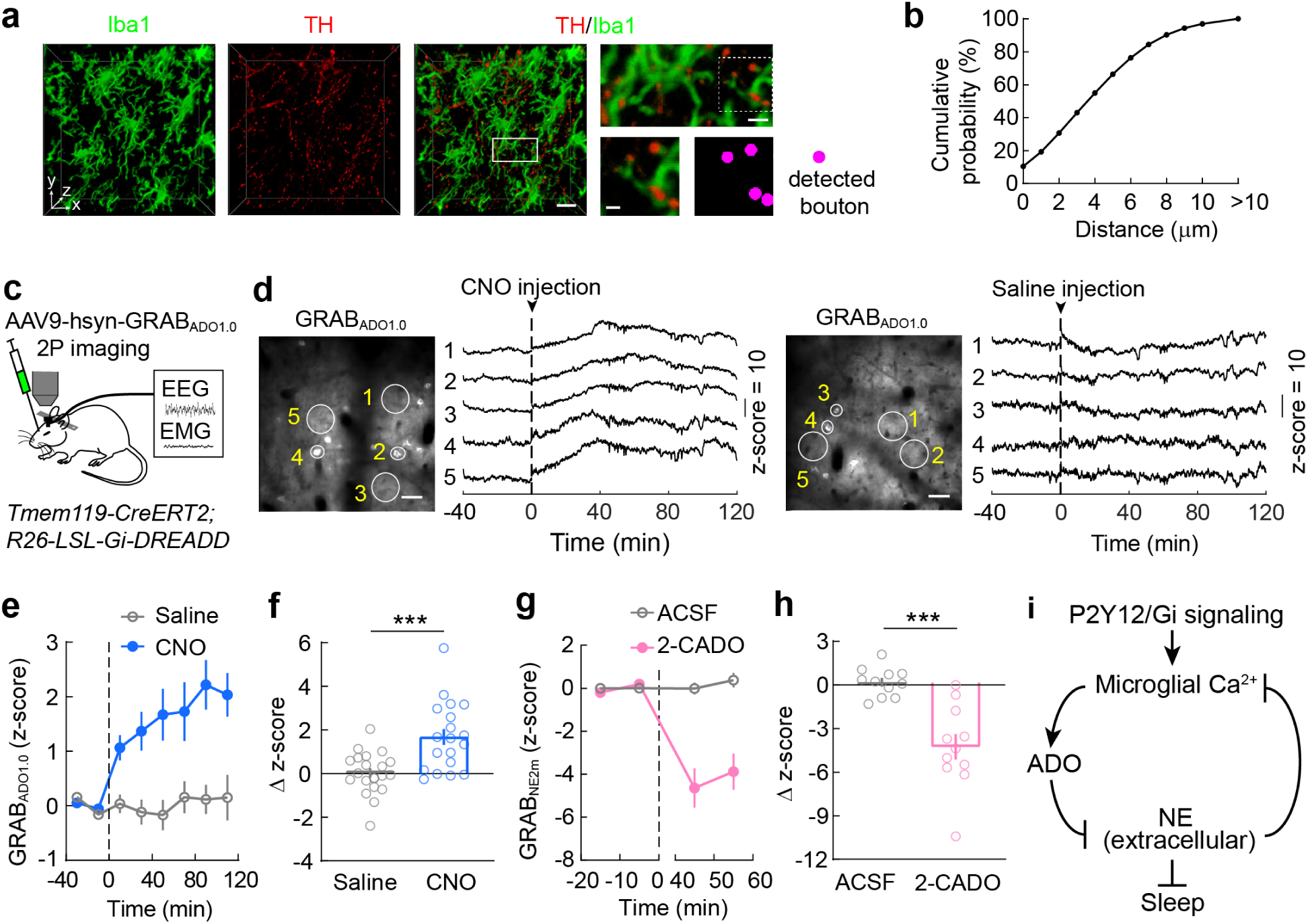
Suppression of NE transmission by microglia Gi signaling is partly mediated by elevated ADO level. **a**, Example images of Iba1-labeled microglia (green) and TH-labeled axons (red) in prefrontal cortex. Left and middle, 3D rendering images of a 50-μm-thick slice; scale bar, 20 μm; top right, high-magnification view of the boxed region; scale bar, 5 μm; bottom right, further enlarged view of the region in dashed box and automatically detected axon buttons (magenta); scale bar, 2 μm. **b**, Distance of buttons to the nearest microglia (n = 140,884 boutons from the prefrontal cortex of 3 mice). **c**, Schematic of GRAB_ADO_ imaging in the prefrontal cortex. **d**, Example imaging session with chemogenetic Gi activation in microglia. Left, field of view (scale bar, 50 μm); 5 ROIs are outlined whose ADO traces are shown on the right. Dashed line, time of CNO or saline injection. **e**, Effect of chemogenetic Gi activation in microglia on ADO signals averaged across 20 (saline) or 19 (CNO) sessions from 6 mice. Dashed line indicates time of injection. **f**, Difference in ADO before and after saline or CNO injection. Each circle represents data from one session. Bars, mean ± s.e.m.; \*\*\**P* < 0.001 (unpaired two tailed *t*-test). **g**, **h**, Extracellular NE levels before and after application of 2-CADO (metabolically stable analog of ADO). Dashed line indicates time of application. Each circle indicates data from one session. 2-CADO, n = 11 sessions; ACSF, n = 11, from 4 mice. Bars: mean ± s.e.m.; \*\*\**P* < 0.001 (unpaired two tailed *t*-test; Δz score was computed as the difference between the periods [-20 0] and [40 60] min). **i**, Diagram summarizing microglia regulation of sleep through reciprocal interactions between microglia Ca^2+^ signaling and NE transmission.

One of the most important wake-promoting neuromodulators is norepinephrine (NE)^36, 37^, which also powerfully modulates microglia motility^38, 39^. We wondered whether NE plays a role in regulating microglia Ca^2+^ activity across brain states. Imaging in the prefrontal cortex of mice expressing a GPCR-activation-based fluorescent NE sensor (GRAB_NE_)^40^ showed a significant increase of cortical NE level at NREM→wake transitions and decrease at wake→NREM transitions (Fig. 4d-f; Extended Data Fig. 6c), although even during NREM sleep we observed NE level fluctuations. These observations are consistent with recent studies based on fiber photometry imaging^41, 42^ as well as previous electrophysiological recordings from locus coeruleus neurons^43^, the main source of NE in the mammalian forebrain. However, they are opposite to the changes in microglia Ca^2+^ activity (Fig. 4a-c).

To test whether NE modulates microglia Ca^2+^, we applied adrenergic receptor antagonists, which can increase NREM sleep (Extended Data Fig. 6d-f). Local application of either ICl-118,551 (ICl, 30 μM, a selective β2 receptor antagonist) or a combination of phentolamine (Phen, 50 μM, antagonist to α receptors) and propranolol (Prop, 10 μM, antagonist to β receptors) in the prefrontal cortex caused a marked increase in microglia Ca^2+^ (Fig. 4g-k), indicating that the reduction of NE signaling during sleep is at least partly responsible for the increased Ca^2+^ activity (Fig. 3a-c). Note, however, that the effect of these antagonists could be mediated either directly by adrenergic receptors on microglia or indirectly through their effects on neurons and astrocytes, which may in turn affect microglia Ca^2+^ activity.

### Activation of microglia Gi suppresses NE transmission

Knowing the effect of NE on microglia Ca^2+^, we wondered whether microglia Ca^2+^ signaling can in turn regulate NE transmission. In *Tmem119-CreERT2; R26-LSL-Gi-DREADD* mice expressing GRAB_NE_ in the prefrontal cortex (Fig. 5a), CNO-induced Gi activation in microglia caused a strong reduction of cortical extracellular NE concentration (Fig. 5b-d; Extended Data Fig. 6g), indicating a mutually antagonistic relationship between NE transmission and microglia Ca^2+^ signaling (Fig. 6i). Given the powerful role of NE in promoting wakefulness, the sleep-promoting effect of microglia P2Y12/Gi signaling is likely mediated at least in part by the reduction of extracellular NE concentration.

To test whether manipulation of microglia within the prefrontal cortex is sufficient for NE reduction, we performed local perfusion of P2Y12 receptor agonist or antagonist. Application of the agonist 2MeSADP (10 μM), which increased microglia Ca^2+^ (Fig. 2g-k), also caused a strong decrease in NE signals, whereas the antagonist PSB0739 (250 μM) caused an NE increase (Fig. 5e-h; Extended Data Fig. 6h). Given the large physical separation between the prefrontal cortex and locus coeruleus, this suggests that microglia can regulate either the release or re-uptake of NE at the axon terminals of locus coeruleus neurons independent of spiking activity of the cell bodies. Indeed, light sheet imaging of tyrosine hydroxylase (TH)-positive axon terminals together with Iba1-labeled microglia confirmed their spatial proximity (Fig. 6a, b; Extended Data Fig. 7a, b), providing ample opportunities for local interactions.

Adenosine (ADO) is known to inhibit the release of neurotransmitters and neuromodulators including NE^44^, and microglia can catabolize ATP/ADP to ADO by the ectonucleotidases CD39 and CD73^21^. We next tested whether activation of microglia Gi signaling affects the extracellular ADO concentration. Imaging in the prefrontal cortex of *Tmem119-CreERT2; R26-LSL-Gi-DREADD* mice expressing the ADO sensor GRAB_ADO_^45^ showed that CNO-induced Gi activation in microglia caused a strong increase in ADO level compared to the control (Fig. 6c-f; Extended Data Fig. 7c). Furthermore, application of 2-Chloroadenosine (2-CADO, 1 μM), a metabolically stable analog of ADO, caused a strong decrease in the NE level (Fig. 6g, h). Together, these results suggest that the suppression of NE transmission by microglia Gi activation is at least partly mediated by an increase in the ADO level (Fig. 6i).

## Discussion

We have shown that activation of Gi signaling in microglia promotes sleep (Fig. 1), and the effect is mediated at least partly by their intracellular Ca^2+^ signaling (Fig. 2, 3), leading to a reduction of extracellular NE concentration (Fig. 5, 6). Although our imaging experiments were performed in the prefrontal cortex (for easy accessibility), region-specific Gi activation through local CNO infusion (which may affect brain tissues within hundreds of microns of the infusion site^46^), suggests that multiple brain regions contribute to the sleep-promoting effect, including the basal forebrain (Extended Data Fig. 7d-r). Given the spatial heterogeneity of microglia^8, 9^, it would be important for future studies to characterize their Ca^2+^ activity in multiple brain areas. Previous studies showed that microglia depletion either increases NREM sleep specifically in the dark phase^15, 16^ or has no significant effect on sleep^17^. A major difference between our study and these earlier studies lies in the time scale: whereas our study focused on dynamic changes over minutes to hours, loss of microglia over days to weeks is likely to cause compensatory changes that may affect sleep through different mechanisms. In future studies it would also be interesting to test the effects of microglia depletion on NE and ADO activity in the brain. Besides microglia, astrocytes can also promote NREM sleep^47–49^. Unlike microglia, however, astrocyte Ca^2+^ activity is higher during wakefulness than both anesthesia^50^ and sleep^47–49^, and it is elevated by NE^51^. One possibility is that the strong elevation of astrocyte Ca^2+^ during prolonged wakefulness^49^ can cause an increase in Ca^2+^-dependent ATP release^52^, which may activate microglia P2Y12/Gi signaling to drive sleep.

P2Y12/Gi signaling is crucial for directing microglia extension towards active neurons^20, 22, 23^ and downregulation of neuronal activity^11, 21, 53, 54^. Here we have shown that Gi activation suppressed NE release (Fig. 5; Extended Data Fig. 6g, h), which may be mediated by increased extracellular adenosine (Fig. 6c-h). In addition to increasing adenosine, microglia also exhibit Ca^2+^-dependent release of cytokines such as tumor necrosis factor alpha (TNFα)^55^, which is known to promote NREM sleep^19^. Interestingly, low doses of TNFα have been shown to promote NREM sleep without affecting REM sleep, but high doses of TNFα also suppressed REM sleep^56^. In our study, Gq activation with a high dose (1 mg/kg) but not low dose (0.2 mg/kg) of CNO caused a suppression of REM sleep (Extended Data Fig. 5f-j), perhaps because at the high CNO dosage Gq signaling induced stronger cytokine release^57^. In addition to P2Y12, 2MeSADP may also activate P2Y1 and P2Y13. In future studies it would be interesting to test the roles of these receptors in brain state regulation.

Chronic sleep restriction can induce both morphological and molecular changes in microglia^58^. Recent studies have shown that during wakefulness microglia exhibit reduced motility compared to anesthetized states, likely due to suppression by NE^38, 39, 59^. Here we show that microglia intracellular Ca^2+^ level changes with brain states (Fig. 4c), which is also mediated at least in part by changed extracellular NE concentration (Fig. 4h-k). In addition to being suppressed by NE (Fig. 4), we showed that microglia Ca^2+^ signaling can in turn cause a rapid reduction of NE (Fig. 5), thus demonstrating a mutually antagonistic relationship between microglia Ca^2+^ and NE transmission in the brain.

Sleep disruption is increasingly recognized as an important risk factor for Alzheimer’s and other neurodegenerative diseases^14^, and the loss of microglia homeostatic functions is associated with both sleep/wake disruption^15, 16^ and disease progression^12, 13^. Our findings point to a mechanistic explanation: an increase in microglia Ca^2+^ enabled by sleep may allow more efficient surveillance and clearance of harmful extracellular proteins involved in neurodegeneration^3,4^; reciprocally, microglia also actively promote sleep for the maintenance of brain homeostasis.

## Supporting information

Legend for Supplementary Video 1

## Methods

### Animals

The following mice were obtained from Jackson Laboratory (Jackson stock number in parenthesis): *Tmem119-CreERT2* (031820), *RCL-GCaMP6s* (028866), *R26-LSL-Gi-DREADD* (026219), *R26-LSL-Gq-DREADD* (026220). All the experiments were performed on adult mice (2 - 6 month) of both sexes. Mice of specific genotype were randomly assigned to experimental and control groups. Experimental and control mice were subjected to exactly the same surgical and behavioral manipulations. Mice were housed in 12 h light-dark cycle (lights on 07:00 and off at 19:00) with free access to food and water. All procedures were approved by Animal Care and Use Committees of the University of California, Berkeley and were done in accordance with federal regulations and guidelines on animal experimentation.

To generate mice with microglia-specific expression of Gi-DREADD and GCaMP6s, we first bred *Tmem119-CreERT2* mice with *R26-LSL-Gi-DREADD*, and then bred the *Tmem119-CreERT2; R26-LSL-Gi-DREADD* mice with *RCL-GCaMP6s* mice, resulting in *Tmem119-CreERT2; RCL-GCaMP6s*; *R26-LSL-Gi-DREADD* mice. A primer set targeting GCaMP6s was used to double check the mouse genotype (5’-AGGACGACGGCAACTACAAG - 3’ and 5’-CACCGTCGGCATCTACTTCA - 3’). Gq-DREADD mice were generated by breeding the *Tmem119-CreERT2* mice with *R26-LSL-Gq-DREADD* mice. To activate tamoxifen-inducible CreERT2, mice were gavaged at ∼6 weeks of age with two doses of 250 mg/kg tamoxifen (T5648, Sigma) in sunflower seed oil (S5007, Sigma) with a separation of 48 h between doses.

### Virus preparation

AAV9-hsyn-GRAB_NE2m_ and AAV9-hsyn-GRAB_Ado1.0_ were obtained from WZ Biosciences Inc. To construct the pAAV-Ef1α-DIO-p130PH-mCherry and pAAV-Ef1α-DIO-p130PH^R134L^-mCherry viral vectors, the transgene p130PH or p130PH^R134L^ were amplified using PCR from the pEGFP-N1 plasmids containing these genes^31^ and inserted into the Ef1α-DIO-mCherry viral vector (Addgene, #47636). AAVTM6 viruses were produced at Janelia Viral Tools facility, and the titer was > 7 × 10^12^ gc/mL. The titer of the NE and ADO sensor viruses was estimated to be ≥ 1 × 10^13^ gc/mL.

### Surgery

For all the surgeries, adult mice were anesthetized with 1.5%–2% isoflurane and placed on a stereotaxic frame. Heating pad was used to keep the body temperature stable during the whole procedure. Eye ointment was applied to keep the eyes from drying. After shaving hairs and asepsis with Betadine and medical alcohol, an incision was made to the skin to expose the skull.

For AAV injection, a craniotomy was made on top of the target region, and AAVs were injected into the target region using Nanoject II (Drummond Scientific) via a micro pipette. 2 μL of AAVTM6-Ef1α-DIO-p130PH-mCherry or AAVTM6-Ef1α-DIO-p130PH^R134L^-mCherry were injected into the lateral ventricle at the coordinates of anteroposterior (AP) +0.1 mm, mediolateral (ML) 0.9 mm, and dorsoventral (DV) 2.5 ∼ 2.8 mm. To activate p130PH or p130PH^R134L^ expression, mice were provided tamoxifen in chow for a two-week period, starting at day 3 post virus injection (250 mg tamoxifen per kg of chow, Research Diets). Mice were gavaged with two additional doses of tamoxifen (250 mg/kg body weight) at days 7 and 9 after virus injection. For the NE and ADO sensors, 0.25 μL of AAV9-hsyn-GRAB_NE2m_ or AAV9-hsyn-GRAB_Ado1.0_ was injected at each of two locations of the prefrontal cortex within the area 0.5 mm ∼ 1.2 mm from the midline and anterior to the bregma, at a depth of 0.5 mm.

For implantation of electroencephalogram (EEG) and electromyogram (EMG) recording electrodes, two miniature stainless-steel screws were inserted into the skull at anteroposterior (AP) −1 mm, mediolateral (ML) 1.5 mm and AP −3 mm, ML 2.5mm. Two EMG electrodes were inserted into the neck musculature. A reference screw was inserted into the skull on top of the right cerebellum. Insulated leads from the EEG and EMG electrodes were soldered to a pin header, which was secured to the skull using dental cement.

For intracerebroventricular and brain-region specific drug infusion, a bilateral cannula (Plastics One Technologies) was inserted into the lateral ventricle (AP +0.1 mm, ML 0.9 mm, and DV 2.6 mm), the prefrontal cortex (AP +1.2 mm, ML 0.5 mm, and DV 0.3 mm), the basal forebrain (AP +0.1 mm, ML 1.2 mm, and DV 5.2mm), and dorsal striatum (AP +0.1 mm, ML 1.5 mm, and DV 2.8 mm). The cannula was placed in the same surgery as EEM and EMG electrode implantation, and was secured to the skull with dental cement.

For cranial window surgery, a dental drill (FST) with 0.5 mm diameter was used to drill through the skull for the craniotomy (3 x 3 mm) over prefrontal cortex. The craniotomy center was ∼1 mm anterior to bregma and 1 mm ML, mostly over the secondary motor area. A 4 x 4-mm glass coverslip (Warner Instruments) was cut with a diamond-point pen and attached to the 3 x 3-mm coverslip by UV glue (Norland Optical Adhesive, Norland). After being sterilized and dried, the combined coverslips were slowly lowered into the craniotomy. A cannula (Plastics One Technologies) was implanted carefully under the cranial window for local drug perfusion.

Dental cement was applied around the window to cover the rim of the glass window. EEG and EMG electrodes were implanted as described above on the opposite side to the cranial window. A stainless-steel head-bar was then firmly attached to the skull and all the remaining exposed skull surfaces were covered by dental cement. Mice were allowed to recover from anesthesia on a heating pad before returned to their home cage. Meloxicam was provided as an analgesic for 24 h post-surgery. In mice with AAV injections, windows were implanted over the injection site after a more than 2-week recovery period.

### Sleep recording

Behavioral experiments were carried out in home cages placed in sound-attenuating boxes between 1:00 pm and 7:00 pm (with CNO injection performed between 1:00 and 2:00 pm and recording completed before 7:00 pm), except for those specifically tested in the dark phase (Extended Data Fig. 2). EEG and EMG electrodes were connected to flexible recording cables via a mini-connector. Recordings started after 20-30 min of habituation. The signals were recorded with a TDT RZ5 amplifier, filtered (0–300 Hz) and digitized at 1,500 Hz. Spectral analysis was carried out using fast Fourier transform (FFT), and brain states were classified into NREM, REM and wake states (wake: desynchronized EEG and high EMG activity, NREM: synchronized EEG with high-amplitude, low-frequency (0.5-4 Hz) activity and low EMG activity, REM: high power at theta frequencies (6-9 Hz) and low EMG activity). The classification was made semi-automatically using a custom-written graphical user interface (programmed in MATLAB, MathWorks).

### Chemogenetic and pharmacological manipulation

For chemogenetic manipulation, saline (0.9% NaCl) or CNO (C0832, Sigma, dissolved in saline) was injected intraperitoneally (i.p.) into mice expressing Gi-DREADD or Gq-DREADD, or mice without DREADD expression. CNO dose was 1 mg/kg body weight for Gi-DREADD or control experiments, and 0.2 mg/kg or 1 mg/kg body weight for Gq-DREADD experiments. Each recording session started immediately after injection and lasted for 5 hours. Each mouse was recorded for 6-8 sessions, and CNO was given randomly in half of the sessions and saline in the other half. Data were averaged across all sessions. For local chemogenetic manipulation, CNO (10 μM, 200 nl, dissolved in ACSF) or ACSF was infused into the target region through bilateral cannula, at 50 nl/min using a microinfusion pump. Each mouse was recorded for 6-8 sessions (CNO was given randomly in half of the sessions and ACSF in the other half) with an interval of at least 2 days. For i.c.v. drug infusion, PSB (1 mM, 2 μl), 2MeSADP (300 μM, 2 μl), ICl (300 μM, 2 ul), a combination of Phen and Prop (Phen:500 μM; Prop: 100 μM; 2 μl) or ACSF was infused into the lateral ventricle through cannula at 500 nl/min.

### Two-photon imaging

Mice with a head-bar were first habituated to sleep under head-fixed condition for two-photon imaging. To do this, the mice were kept head-fixed under the two-photon system for ∼ 15 min, ∼ 30 min, and ∼ 45 min for the first three days. The duration of head-fixation increased by 20 to 30 min in each subsequent session, reaching a maximum of ∼ 3 h. EEG and EMG signals were recorded during later sessions to monitor the state of the mouse until multiple wake-sleep cycles were observed.

During imaging sessions, the mouse was allowed about 10 min of habituation after being head-fixed before imaging started. For imaging with chemogenetic treatment, a 40 min baseline period was imaged before saline or CNO injection, and another 80 to 120 min was imaged after injection. For imaging with drug application, a 20 min baseline period was imaged before drug perfusion. Drug was perfused to the cortical surface through an infusion pump (Micro4, World Precision Instruments) at a rate of 0.5 μL/min (2 μL volume in total). The following drugs were used: ICl-118,551 (I127, Sigma), Phentolamine Mesylate (6431, Tocris), Propranolol (P0884, Sigma), PSB0739 (3983, Tocris), 2MeSADP (1624/10, R&D), and 2-CADO (C5134, Sigma). All the drugs are constituted with ACSF (3525, Tocris).

Two-photon Ca^2+^ imaging, GRAB_NE2m_ or GRAB_ADO1.0_ imaging was performed using a custom two-photon microscope that has been described previously^60^. EEG and EMG were recorded simultaneously with a TDT RZ5 amplifier as described above. The microscope (Movable Objective Microscope; Sutter Instrument) was controlled by the ScanImage software and the objective was a 20× water immersion lens (XLUMPlanFI, 0.95 NA; Olympus). A Mai-Tai Insight laser (Spectra-Physics) was tuned to 920nm and ∼ 35 mV output for microglia GCaMP6s imaging, < 10 mV output for GRAB_NE2m_ and GRAB_ADO1.0_ imaging (as measured under the objective). Fluorescence emission was collected using a GaAsP PMT (H10770PA-40; Hamamatsu). Microglia Ca^2+^ was imaged at a frame rate of 0.84 Hz, 512 × 512 pixels, and 2x digital zoom. NE and ADO signals were imaged at a frame rate of 1.68 Hz and pixel resolution of 256 × 256.

### Calcium imaging data analysis

Time series of Ca^2+^ activity images were motion corrected with Inscopix Data Processing software. An average intensity image was then generated for region of interest (ROI) selection. Microglia with clear soma and processes was semi-automatically tracked with Simple Neurite Tracer plugin in ImageJ. The morphology mask of the cell was obtained by applying the ‘Fill’ function within the plugin. Microglia soma and processes were then manually segmented into microdomains using the ‘freehand selections’ tool in ImageJ. Soma was identified based on the size and the intensity. Processes were segmented if greater than 5 μm. The mean intensity values were generated with the multi-measure tool in ImageJ for the segmented ROIs. The mean fluorescence intensity and the standard deviation during the baseline period was used to obtain the z-score of GCaMP6s signal. For the amplitude and frequency analysis, we adopted a previously described method^30^. Briefly, the baseline fluorescence of the ROI, F_0_, was determined as the lower 25^th^ quartile of the fluorescence in a ± 600-s sliding window and was used to calculate ΔF/F. A calcium transient was considered to occur at a threshold that is 3x standard deviation above the mean value of the ΔF/F trace over 3 frames.

### GRAB_NE2m_ and GRAB_ADO1.0_ imaging data analysis

Time series of GRAB_NE2m_ or GRAB_ADO1.0_ images were motion corrected by a custom MATLAB script using an open-source toolkit ANTs (picsl.upenn.edu/software/ants/). An average intensity image was then generated for ROI selection. Cell bodies were identified based on the intensity. In regions without clear cell bodies, ROIs with diameters of 50 − 70 μm were selected. 8 to 12 ROIs were manually selected across the image. The mean intensity of each ROI was generated over time. The mean fluorescence intensity and the standard deviation during the baseline period was used to obtain the z-score of GRAB_NE2m_ or GRAB_ADO1.0_ signals.

### Microglia-bouton distance analysis

For the quantification of the distance between boutons and microglia, wildtype brain or brain with AAV2-EF1α-DIO-eYFP (250 nl) injected into the locus coeruleus (AP −5.4 mm, ML 0.9 mm, DV 3.7 mm) of *Dbh-Cre* mouse was perfused using PBS followed by 4% paraformaldehyde in PBS. Brains were post-fixed in 4% paraformaldehyde for 24 h. Further tissue processing, immunolabeling (TH-antibodies, Iba1 antibodies, GFP antibodies), light sheet imaging of the prefrontal cortex, and image registration were done by LifeCanvas Technologies with SmartSPIM at 1 μm z-step and 0.41 μm x-y pixel size. Regions with relatively even distribution of labeled axons were manually selected. Automatic denoising (with the same parameters), manual adjustment of contrast and brightness were used as pre-processing step to improve image quality through the ImageJ software (NIH). Subsequently, DeepBouton^61^ was used to detect the boutons. An iterative weakly supervised segmentation network was also used to extract the microglia from the Iba1 images, which is an improved version of 3D Res-UNet^62^. Finally, the identified boutons were verified manually and the minimal distance from axon bouton to microglia was calculated. The 3D rendering of the example images in Fig. 6a and Extended Data Fig. 7a were generated with Imaris software (BITPLANE).

### Immunohistochemistry

For immunohistochemistry, mice were deeply anesthetized and transcardially perfused using PBS followed by 4% paraformaldehyde in PBS. Brains were post-fixed in 4% paraformaldehyde for 24 - 48 h and stored in 30% sucrose in PBS solution for 48 h for cryoprotection. Brains were embedded and mounted with Tissue-Tek OCT compound (Sakura finetek) and 30 μm sections were cut using a cryostat (Leica). For P2Y12 staining, brain slices were washed using PBS and subjected to heat-induced antigen retrieval with 0.1 M citric acid (pH 6.0) at 95°C for 5 min.

Slices were then washed with PBS, permeabilized using PBST (0.3% Triton X-100 in PBS) for 30 min and incubated with blocking solution (5% normal bovine serum in PBST) for 1 h followed by rabbit anti-P2Y12 antibody (AS-55043A, AnaSpec) together with goat anti-Iba1 antibody (ab5076, Abcam) incubation overnight at 4 °C. The next day, after sufficient washing in PBS, sections were incubated in proper fluorescently conjugated secondary antibodies (1:500, Invitrogen) for 2 h at room temperature. Finally, sections were counterstained with 4’,6-diamidino-2-phenylindole dihydrochloride (DAPI; Sigma-Aldrich) and mounted on slides with VECTASHIELD Antifade Mounting Medium (Vector Laboratories, H-1000).

For HA staining, Alexa Fluor 488 Tyramide SuperBoost Kit (B40922, Thermo Fisher Scientific) was used. Briefly, floating brain slices were treated with 3% hydrogen peroxide solution for 15 min to quench the endogenous peroxidase activity and then subjected to heat-induced antigen retrieval as described above. After wash with PBS, slices were incubated in blocking buffer (5% normal goat serum) for 1 h. Rabbit anti-HA antibodies (#3724, Cell Signaling Technology) together with chicken anti-Iba1 (#234009, Synaptic Systems) were applied overnight at 4°C. Afterward, sections were washed three times with PBS, 10 min each. Alexa Fluor 647 anti-chicken IgG together with Poly-HRP-conjugated anti-rabbit secondary antibody was applied to the slices for 2 h at room temperature. Slices were washed with PBS three times, 10 min each. Tyramide working solution was prepared according to the manufacturer’s instructions, and colors were developed with Alexa Fluor 488 Tyramide for 10 to 30 min. Reaction stop reagent was applied, and slices were counterstained with DAPI and mounted with antifade mounting medium.

For p130PH-mCherry staining, brain slices were washed using PBS and permeabilized using PBST for 30 min and incubated with blocking solution (5% normal goat serum in PBST) for 1 h. Rat anti-mCherry antibody (M11217, Life Technologies) together with chicken anti-Iba1 and rabbit anti-P2Y12 antibody were applied overnight at 4 °C. The next day, after sufficient washing in PBS, sections were incubated with Biotin-SP-conjugated anti-rat (1:500, Jackson ImmunoResearch), Alexa Fluor 647 anti-chicken, and Alexa Fluor 488 anti-rabbit secondary antibodies for 2 h at room temperature. Sections were sufficiently washed again, and Alexa Fluor 594-conjugated streptavidin (1:1000, Jackson ImmunoResearch) was applied to the slices for 1.5 h at room temperature. Afterward, slices were counterstained with DAPI and mounted with antifade mounting medium.

Fluorescence images were taken with a confocal microscope (LSM 710 AxioObserver Inverted 34-Channel Confocal, Zeiss, at 20x, for Figs.1 and 3, and Extended Data Figs. 1 and 4) or a fluorescence microscope (Keyence BZX-710, for Extended Data Figs. 6c and 7c, e, j, o). Cell counting was done with custom-written MATLAB software in a ∼1000 μm × 500 μm - 1000 μm × 1000 μm of region in each brain area.

### Statistics

Statistical analysis was performed using GraphPad Prism, and significance was determined at *P* < 0.05. All statistical tests were two-sided. The selection of statistical tests was based on reported previous studies. A normality test was performed on each dataset using the Shapiro-Wilk test or D’Agostino & Pearson test. For comparison of two group means, the parametric tests (paired *t* test or unpaired *t* test) were used if the dataset was normally distributed; otherwise, nonparametric tests (Wilcoxon signed rank test or Mann-Whitney *U* test) were used. One-way ANOVA was used for comparison across more than two groups. Two-way ANOVA with Bonferroni correction was used for comparisons of brain state between different conditions (saline vs. CNO) in chemogenetic experiments.

## Data availability

The datasets used for the figures is available upon request from the authors.

## Code availability

Custom-written MATLAB code is available upon request from the authors.

## Acknowledgments

We thank Tamas Balla and Gyorgy Hajnoczky for generously sharing the p130PH and p130PH^R134L^ constructs, Bryan Roth and Peter Grace for sharing information on DREADD tools, Hui Gong and Xiangning Li for the discussion of automated bouton detection, Hongfeng Gao for assistance with obtaining the transgenic mouse lines, Min Xu for providing the program for sleep data analysis. This work was supported by the Howard Hughes Medical Institute.

## Author Contributions

C.M. and Y.D. conceived and designed the study and wrote the paper with inputs from all other authors. C.M. and B.L. performed most of the experiments. B.L. wrote programs for data analysis. C.M. and B.L. analyzed the data. D.S. and C.T. helped obtain and package some of the AAV viruses. D.S., Y.H., and Y.Z. helped with brain state classification. X.D. helped with histological staining. Y.Z. quantified histological images. A.L., C.X., G.H., and Z.X. analyzed the light-sheet imaging data. K.W. and S.M. made significant contribution during the exploratory phase of the project. W.C. helped with 2P light path modifications and corrections. K.S. and Q.L. provided critical consultation and feedback. Y.D. supervised all aspects of the work.

## Additional Information

Supplementary Information is available for this paper. Correspondence and requests for materials should be addressed to Y.D.

---

**Extended Data Fig. 1.**
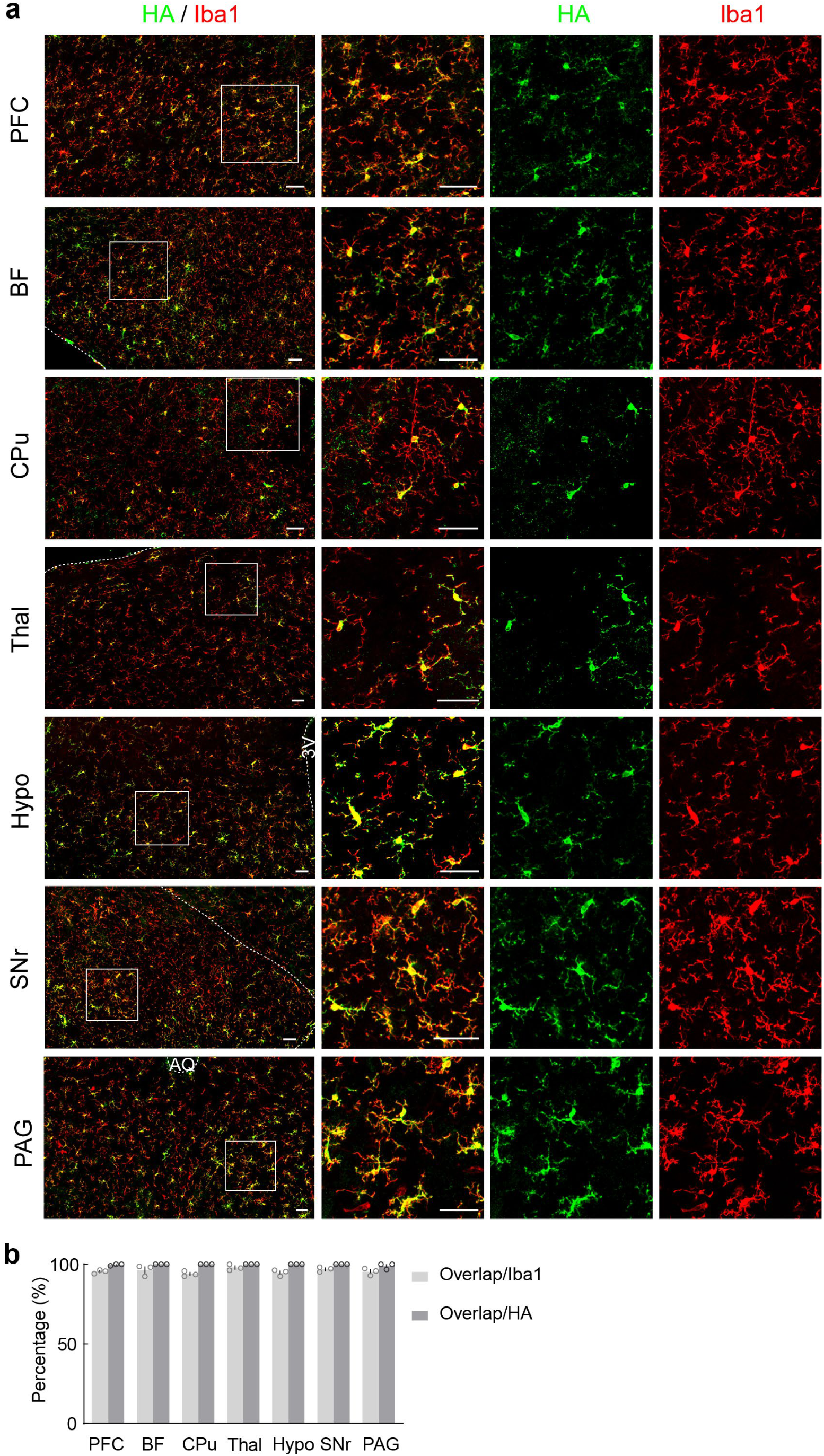
Specificity and efficiency of Gi-DREADD expression in microglia in *Tmem119-CreERT2; R26-LSL-Gi-DREADD* mice. **a**, Confocal images from multiple brain regions showing hM4Di (Gi-DREADD) expression (detected by an HA-tag antibody) in Iba1+ microglia. White box in left panel, region enlarged on the right. Scale bar, 50 μm. PFC, prefrontal cortex; BF, basal forebrain; CPu, caudate putamen; Thal, thalamus; Hypo, hypothalamus; SNr, substantia nigra pars reticulata; PAG, periaqueductal gray. b, Quantification of efficiency and specificity (n = 3 mice).

**Extended Data Fig. 2.**
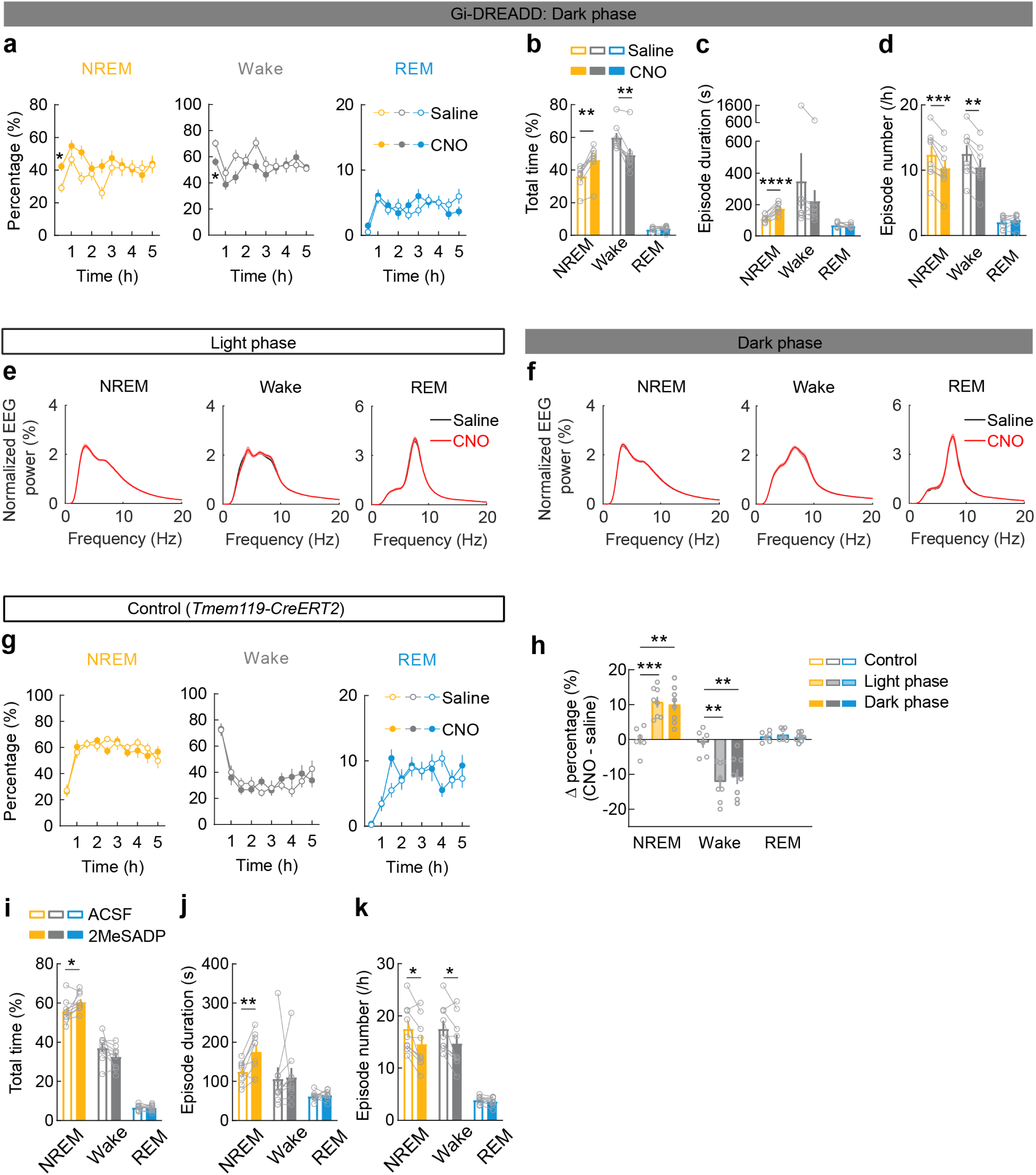
Effect of Gi-DREADD activation on sleep in dark phase, EEG power spectra within each state, control experiments in mice without Gi-DREADD, and effect of P2Y12 agonist on sleep. **a**, Summary of chemogenetic experiment in dark phase (beginning at ∼8:00 pm). Shown are the percentages of time in each brain state following CNO and saline injection (mean ± s.e.m; n = 8 mice; two-way ANOVA with Bonferroni correction; NREM: treatment, *P* = 0.020; time, *P* =0.012; Wake: treatment, *P* = 0.040; time, *P* = 0.0039; REM: treatment, *P* = 0.89; time, *P* < 0.0001). **b**, Percentage of time in each brain state within 3 h after CNO or saline injection. Each circle indicates data from one mouse (mean ± s.e.m; n = 8 mice). \*\**P* < 0.01 (Wilcoxon signed rank test). **c**, **d**, Mean episode duration (**c**) and episode number per hour (**d**) for each brain state within 3 h after CNO or saline injection. Each circle indicates data from one mouse (mean ± s.e.m; n = 8 mice). \*\**P* < 0.01, \*\*\**P* < 0.001, \*\*\*\**P* < 0.0001 (paired two-tailed *t*-test or Wilcoxon signed rank test). **e**, **f**, Comparison of normalized EEG power spectra within each brain state between saline and CNO sessions during the light phase (**e**) or dark phase (**f**), averaged across 8 mice. The EEG spectra of each session was normalized by the total power between 0 and 25 Hz before averaging. **g**, Percentages of time in each brain state following CNO or saline injection in *Tmem119-CreERT2* control mice without Gi-DREADD expression (mean ± s.e.m; n = 6 mice; two-way ANOVA with Bonferroni correction; NREM: treatment, *P* = 0.74; time, *P* < 0.0001; Wake: treatment, *P* = 0.82; time, *P* < 0.0001; REM: treatment, *P* = 0.77; time, *P* < 0.0001). **h**, Changes in each brain state induced by chemogenetic manipulation (difference between CNO and saline injections, averaged across 3-h after injection) in mice without Gi-DREADD (control) or with Gi-DREADD (treatment during light phase or dark phase). Each circle indicates data from one mouse; error bar: ± s.e.m. \*\**P* < 0.01 (One-way ANOVA with Holm-Šídák’s test; NREM, *P* = 0.0004; Wake, *P* = 0.0018; REM, *P* = 0.60). **i-k**, Percentage of time (**i**), mean episode duration (**j**), and episode number per hour (**k**) for each brain state within 2 h after i.c.v. infusion of P2Y12 agonist (2MeSADP) or ACSF. Each circle indicates data from one mouse; error bar: ± s.e.m (n = 9 mice). \**P* < 0.05, \*\**P* < 0.01 (paired two-tailed *t*-test or Wilcoxon signed rank test).

**Extended Data Fig. 3.**
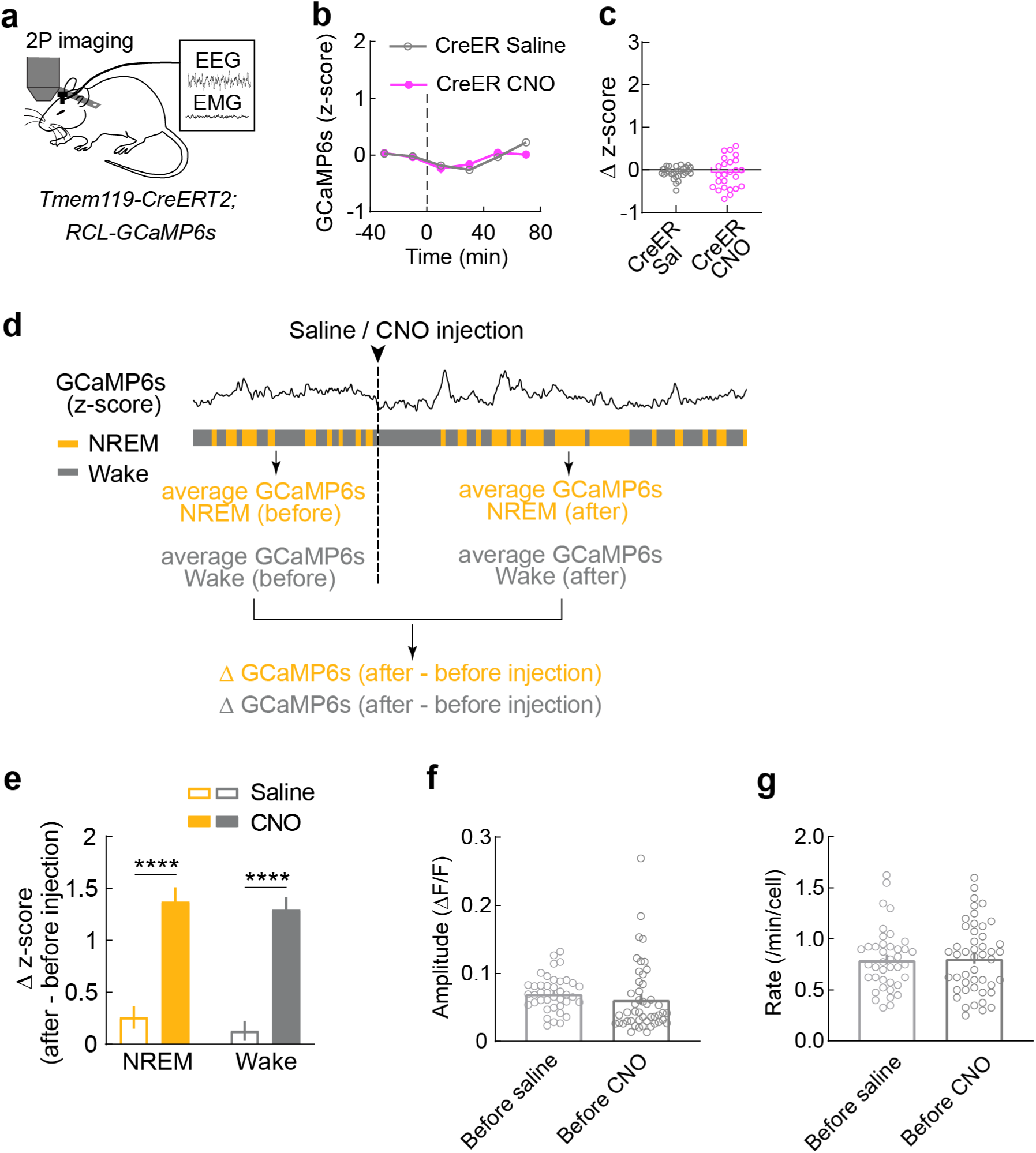
Control experiment for Ca^2+^ imaging in mice without Gi-DREADD, comparison of microglia Ca^2+^ activity before and after CNO-induced Gi activation within each brain state and Ca^2+^ activity during baseline period. **a**, Schematic for two-photon (2P) Ca^2+^ imaging in head-fixed mice without Gi-DREADD expression. **b**, Z-scored Ca^2+^ activity averaged across all microglia (CNO: n = 25; saline: n = 28; from 4 mice). Dashed line, time of injection. **c**, Population summary of the change in mean Ca^2+^ level (*P* =0.73; Mann-Whitney *U* test). **d**, Diagram illustrating the comparison of Ca^2+^ activity before and after CNO or saline injection within each brain state. First, Ca^2+^ activity in NREM or wakeful episodes are averaged separately for before and after injection; second, the difference between the averaged Ca^2+^ activity before and after injection are calculated for each brain state. **e**, CNO-induced change in microglia Ca^2+^ for each brain state (difference between before and after injection). **f**, **g**, Amplitude (**e**) and frequency (**f**) of Ca^2+^ events during baseline period (before saline or CNO injection). CNO: n = 47; saline: n = 40; from 5 mice (amplitude, *P* = 0.30; frequency, *P* = 0.86; unpaired two-tailed *t*-test).

**Extended Data Fig. 4.**
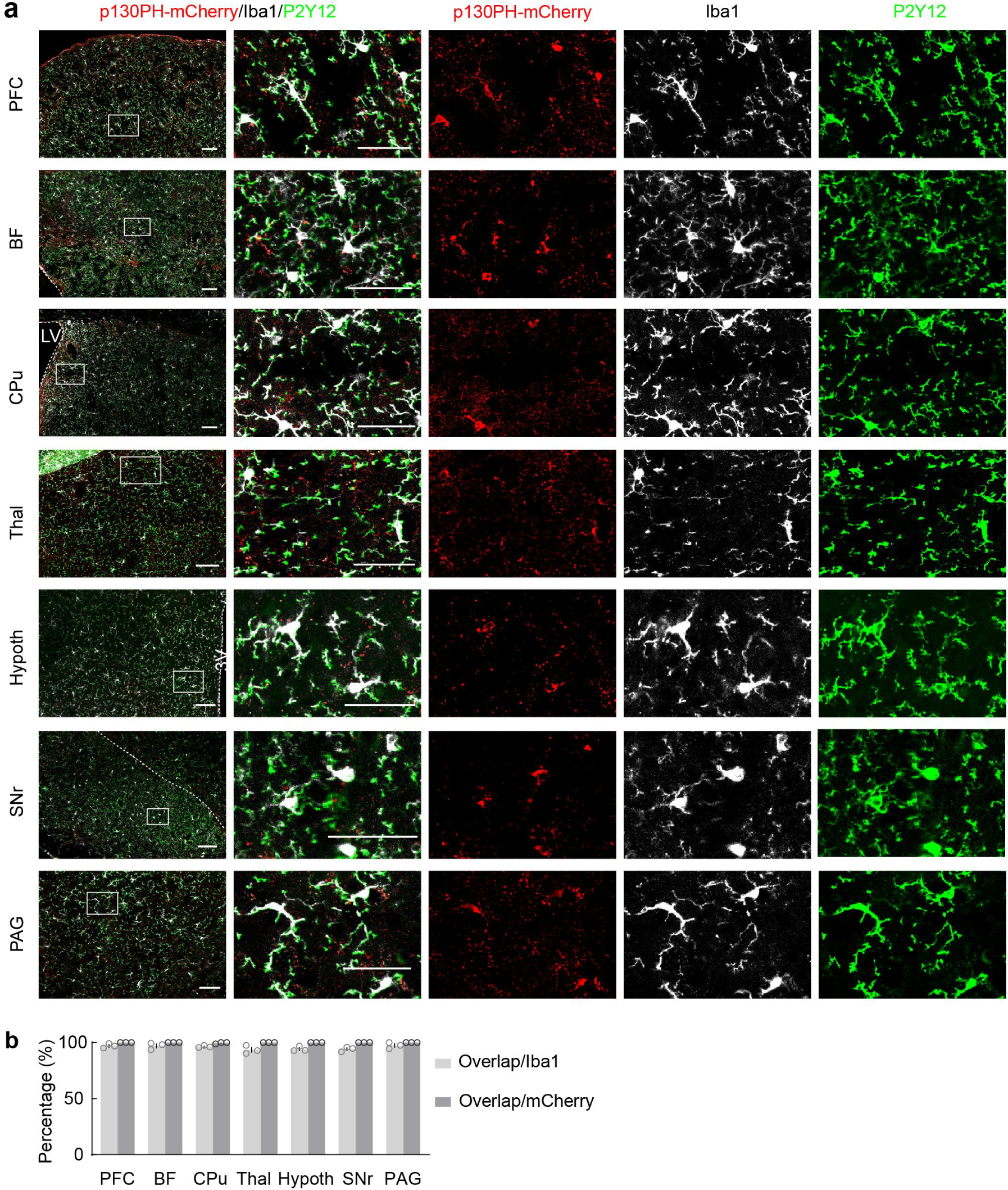
Efficiency and specificity of p130PH-mCherry expression in microglia. **a**, Confocal images of p130PH-mCherry expression in Iba1+, P2Y12+ microglia in multiple brain regions. White box in left panel, region enlarged on the right. Scale bar: left, 100 μm; right, 50 μm. PFC, prefrontal cortex; BF, basal forebrain; CPu, caudate putamen; Thal, thalamus; Hypo, hypothalamus; SNr, substantia nigra pars reticulata; PAG, periaqueductal gray. **b**, Quantification of efficiency and specificity (n = 3 mice).

**Extended Data Fig. 5.**
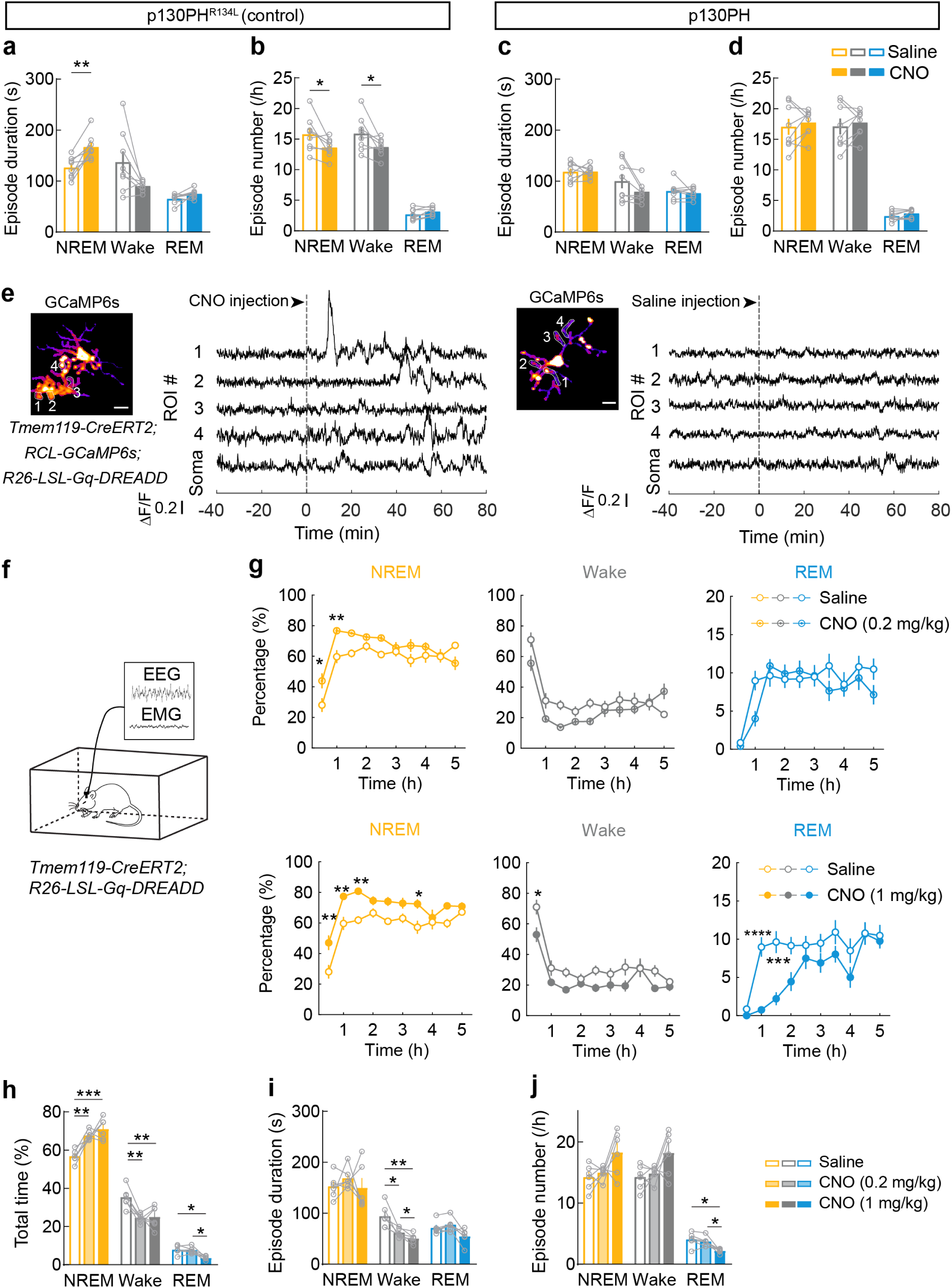
Effects of Gi-DREADD activation in mice expressing p130PH^R134L^ or p130PH and effect of Gq-DREADD activation on microglia Ca^2+^ and sleep. **a, b**, Mean episode duration (**a**) and episode number per hour (**b**) for each brain state within 3 h after CNO or saline injection in mice expressing p130PH^R134L^. Each circle indicates data from one mouse; bars: mean ± s.e.m (n = 8 mice). \**P* < 0.05, \*\**P* < 0.01 (paired two-tailed *t*-test). **c, d**, Similar to (**a, b**), but for mice expressing p130PH (n = 8 mice; **c**, NREM: *P* = 0.94; Wake: *P* = 0.088; REM: *P* = 0.46; **d**, NREM: *P* = 0.52; Wake: *P* = 0.55; REM: *P* = 0.16; paired two-tailed *t*-test). **e**, Example imaging sessions with CNO and saline injection in mice expressing GCaMP6s and Gq-DREADD in microglia. Left, representative microglia (scale bar, 10 μm); 4 ROIs in microglia processes are outlined, whose Ca^2+^ traces are shown on the right. Dashed line indicates time of CNO or saline injection. **f**, Schematic for chemogenetic experiment with Gq-DREADD mice. **g**, Percentages of time in each brain state following CNO or saline injection (top, CNO: 0.2 mg/kg; bottom, CNO: 1 mg/kg; mean ± s.e.m; n = 6 mice). \**P* < 0.05, \*\**P* < 0.01, \*\*\**P* < 0.001, \*\*\*\**P* < 0.0001 (two-way ANOVA with Bonferroni correction; top, NREM: treatment, *P* < 0.0001; time, *P* < 0.0001; Wake: treatment, *P* = 0.0043; time, *P* < 0.0001; REM: treatment, *P* = 0.78; time, *P* < 0.0001; bottom, NREM: treatment, *P* < 0.0001; time, *P* < 0.0001; Wake: treatment, *P* = 0.0002; time, *P* < 0.0001; REM: treatment, *P* = 0.0002; time, *P* < 0.0001). **h-j**, Percentage of time (**h**), mean episode duration (**i**) and episode number per hour (**j**) for each brain state within 3 h after CNO or saline injection. Each circle indicates data from one mouse, bars: mean ± s.e.m (n = 6 mice). \**P* < 0.05, \*\**P* < 0.01, \*\*\**P* < 0.001 (One-way ANOVA with Holm-Šídák’s test; H, NREM, *P* = 0.0029; Wake, *P* = 0.0037; REM, *P* = 0.0011; I, NREM, *P* = 0.55; Wake, *P* = 0.003; REM, *P* = 0.028; J, NREM, *P* = 0.074; Wake, *P* = 0.083; REM, *P* = 0.013).

**Extended Data Fig. 6.**
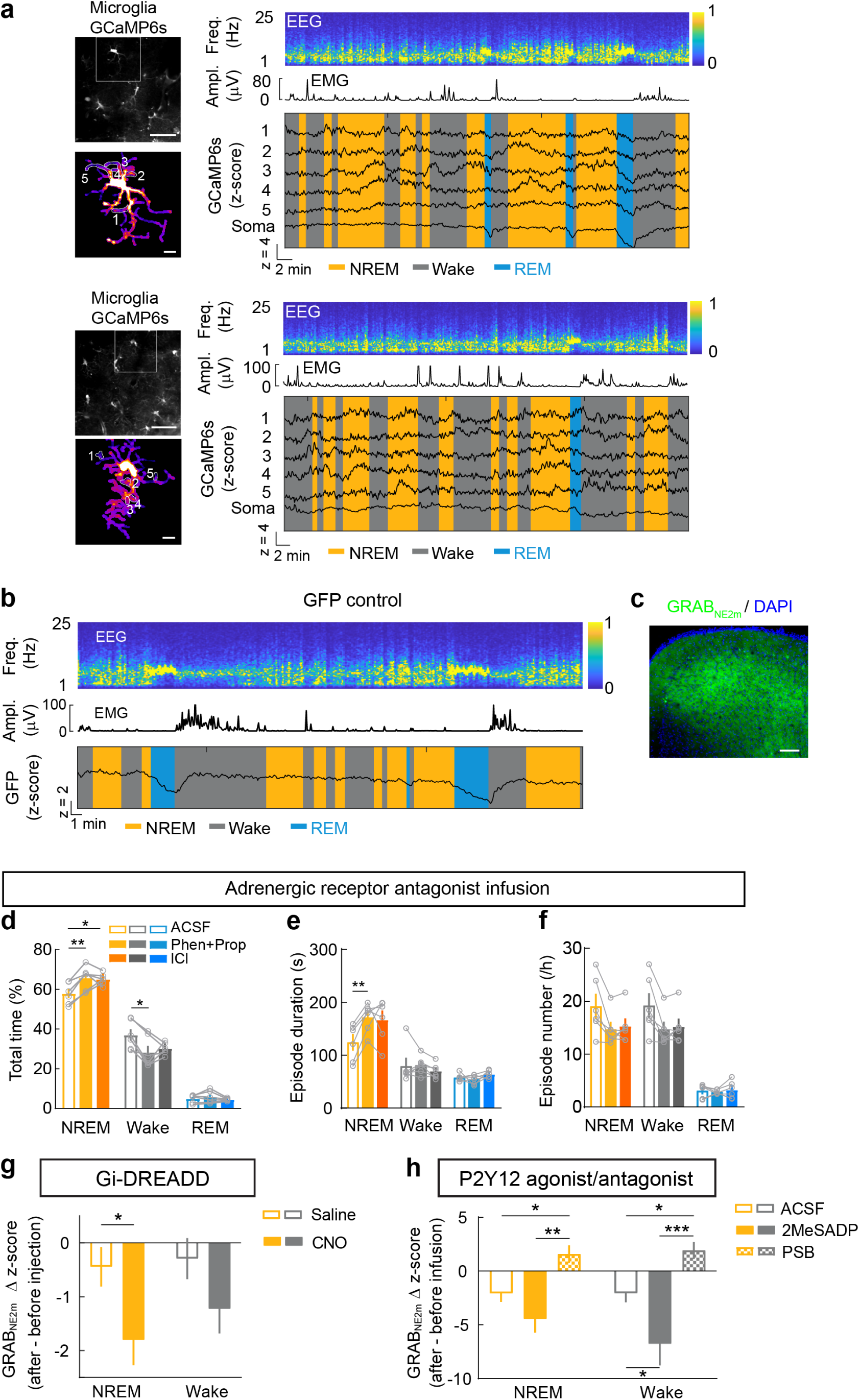
Ca^2+^ imaging sessions with episodes of wakefulness, NREM sleep, and REM sleep, control experiment for 2P imaging with Ca^2+^-independent GFP, GRAB_NE_ expression, effect of adrenergic receptor antagonists on sleep, and comparison of cortical NE activity before and after microglia manipulations within the same brain state. **a**, Two examples of Ca^2+^ imaging session. Top left, field of view containing multiple microglia (scale bar, 50 μm); Bottom left, high magnification view of the microglia soma and processes in the white box (scale bar, 10 μm); 5 ROIs in processes are outlined, whose Ca^2+^ traces are shown on the right together with EEG spectrogram (Freq., frequency), EMG amplitude (Ampl.), and color-coded brain states. **b**, An example 2P imaging session with GFP showing consistent decrease in fluorescence during REM sleep. Top, EEG spectrogram (Freq., frequency); middle, EMG amplitude (Ampl.); bottom, GFP signal (from a *Cx3cr1^eGFP/+^* mouse) and color-coded brain states. **c**, Image showing the expression of the GRAB_NE2m_ in the prefrontal cortex. Scale bar, 100 μm. **d-f**, Percentage of time (**d**), mean episode duration (**e**), and episode number per hour (**f**) within 3 h after i.c.v. infusion of Phen (α receptor antagonist) and Prop (β receptor antagonist), ICl (β2 receptor antagonist), or ACSF. Each circle indicates data from one mouse; bars, mean ± s.e.m; n = 6 mice. \**P* < 0.05, \*\**P* < 0.01 (One-way ANOVA with Holm-Šídák’s test; **d**, NREM, *P* = 0.0058; Wake, *P* = 0.023; REM, *P* = 0.37; **e**, NREM, *P* = 0.037; Wake, *P* = 0.42; REM, *P* = 0.18; **f**, NREM, *P* = 0.053; Wake, *P* = 0.048; REM, *P* = 0.72;). **g**, Change of NE activity in each brain state induced by chemogenetic activation of microglia Gi signaling (difference between before and after injection) (mean ± s.e.m, saline, n = 13 sessions, CNO, n = 14; from 5 mice). \**P* < 0.05 (unpaired two-tailed *t*-test). **h**, Similar to (**g**), but for local perfusion experiments (mean ± s.e.m; 2MeSADP, n = 8 sessions; PSB, n = 7; ACSF, n =10, from 4 mice). \**P* < 0.05, \*\**P* < 0.01, \*\*\**P* < 0.001 (One-way ANOVA with Holm-Šídák’s test; NREM, *P* = 0.002; Wake, *P* = 0.001).

**Extended Data Fig. 7.**
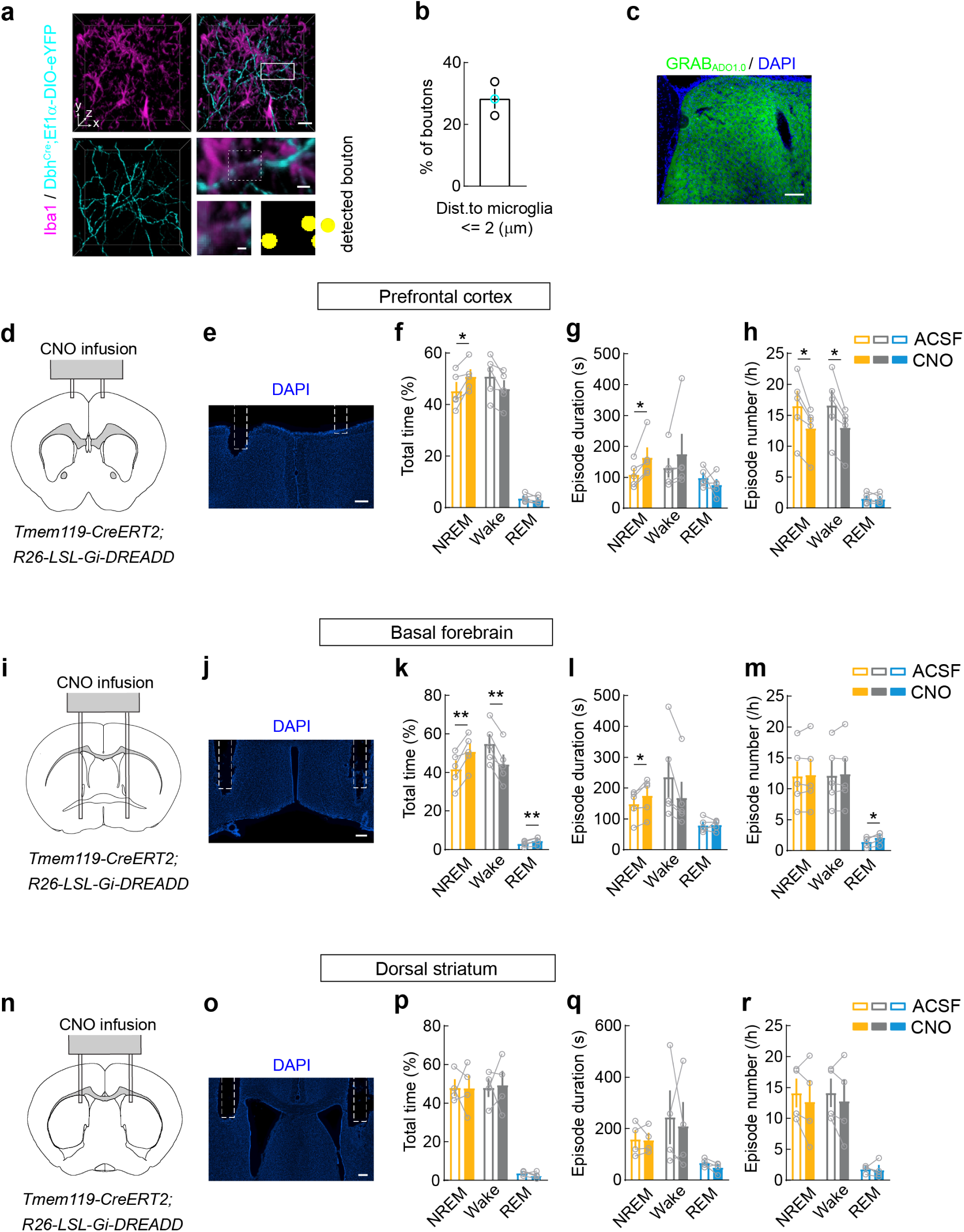
Distance of adrenergic axon boutons to microglia, GRAB_ADO_ expression, and effect of local Gi-DREADD activation on sleep. **a**, Example images of Iba1-labeled microglia (magenta) and Dbh-eYFP-labeled axons (cyan) in prefrontal cortex. Left and top right, 3D rendering images of a 50-μm-thick slice; scale bar, 20 μm; middle right, high-magnification view of the boxed region; scale bar, 5 μm; bottom right, further enlarged view of the region in dashed box and automatically detected axon buttons (yellow); scale bar, 2 μm. **b**, Percentage of buttons within 2 μm from the nearest microglia. Each circle represents data from one mouse (n = 3 mice; black circle, TH-labeled axon boutons detected by immunohistochemistry; cyan circle, Dbh-eYFP-labeled boutons). Dist., distance. **c**, Image showing the expression of the GRAB_ADO_ in the prefrontal cortex. Scale bar, 100 μm. **d, e**, Coronal diagram (**d**) and image (**e**) showing local infusion site in the prefrontal cortex of *Tmem119-CreERT2; R26-LSL-Gi-DREADD* mice. Scale bar, 200 μm. **f-h**, Percentage of time (**f**), mean episode duration (**g**), and episode number per hour (**h**) for each brain state within 2.5 h after CNO or ACSF infusion. Each circle indicates data from one mouse (mean ± s.e.m; n = 5 mice). \**P* < 0.05, \*\**P* < 0.01 (paired two-tailed *t*-test or Wilcoxon signed rank test). **i-m**, Similar to (**d-h**), but for basal forebrain infusion (n = 5 mice). **n-r**, Similar to (**d-h**), but for dorsal striatum infusion (n = 4 mice).

