## Supplementary material for "Microglia Regulate Sleep via Calcium-Dependent Modulation of Norepinephrine Transmission": Legend for Supplementary Video 1

Two-photon imaging of Ca^2+^ activity of a representative microglia before and after CNO-induced Gi-DREADD activation in *Tmem119-CreERT2; RCL-GCaMP6s; R26-LSL-Gi-DREADD* mice, related to data shown in Fig. 2c-f and Extended data Fig. 3.
